# The endogenous metabolite hypochlorite activates indoleamine 2,3-dioxygenase-1 for catalysis: Functional and mechanistic implications

**DOI:** 10.64898/2026.04.21.719948

**Authors:** Setareh Saryazdi, Steven G. Van Lanen

## Abstract

Indolamine 2,3-dioxygenase (IDO1) is a hemoprotein that catalyzes the oxidative cleavage of L-tryptophan (L-Trp) to *N’*-formyl-L-kynurenine (L-NFK) along the kynurenine pathway. Its activity depletes L-Trp while initiating a signaling cascade culminating in an immunosuppressive outcome of clinical significance. The generally used in vitro activity assay for IDO1 relies on ascorbic acid and synthetic methylene blue, with the endogenous activator still uncertain. Here we demonstrate sodium hypochlorite, commonly known as bleach, functions as an in vitro activator/cofactor for the recombinant human IDO1-catalyzed dioxygenation reaction. Other hypohalous acids, generated in situ by lactoperoxidase (LPO) or in aqueous solutions of I_2_ or Br_2_, also activate IDO1. Contrastingly, the pseudohalide thoicyanate, a known, excellent substrate for LPO, yielded only trace levels of L-NFK. Importantly, complete conversion to L-NFK occurs with sub-stoichiometric hypohalous acid relative to L-Trp, and the overall reaction remains O_2_-dependent. Kinetic analysis with variable L-Trp and multiple, fixed concentrations of the activators/ cofactors revealed typical Michaelis-Menten kinetics without substrate inhibition, which contrasts past analysis using alternative IDO1 assays. The calculated second order rate constants were overall comparable regardless of the identity and concentration of hypohalous acid. Finally, 1-methyl-L-tryptophan, a reported inhibitor and poor substrate for rhIDO1, was reexamined with the hypohalous acid-dependent conditions revealing an improved catalytic efficiency when compared with the native substrate L-Trp. Along with this unanticipated result, the in vivo functional and mechanistic implications of the newly discovered hypohalous acid-dependent IDO1 activity are discussed.

**Significance:** Indoleamine 2,3-dioxygenase-1 (IDO1) is a hemoprotein that catalyzes the conversion of L-tryptophan to *N’*-formyl-L-kynurenine (L-NFK). Past efforts have culminated in the conclusion that IDO1 is a checkpoint modulator of mammalian immunity, both in terms of the immunogenicity and immune tolerance. However, the identity of the endogenous activator of IDO1 is still unsettled. Here we report that sodium hypochlorite, aka bleach, activates ferric rhIDO1 in a catalytic manner, enabling rhIDO1 to efficiently catalyze the production of L-NFK in an O_2_-dependent reaction. The results suggest rhIDO1 can efficiently operate via a hypochlorous acid-dependent mechanism that is reminiscent of the peroxide-shunt pathway used by other hemoproteins. Furthermore, the results suggest a functional role for endogenously produced hypochlorous acid beyond solely killing foreign pathogens.

## INTRODUCTION

L-Tryptophan (L-Trp), an essential amino acid in the animal diet, is predominantly metabolized via the kynurenine (Kyn) pathway with estimations of 95% of consumed L-Trp being used in this capacity (*1*). Dysregulation or abnormalities of the Kyn pathway have been implicated in several pathologies including neurological diseases, psychiatric disorders, inflammatory conditions, and cancers, among many others (*2-4*). The first enzymatic step in the Kyn pathway, the oxidative conversion of L-Trp to *N’*-formyl-L-kynurenine (L-NFK), is catalyzed by one of three distantly related isozymes—tryptophan 2,3-dioxygenase (TDO), indoleamine 2,3-dioxygenase-1 (IDO1) or IDO2 (*5*) (Figure 1). Although still the subject of on-going research, the isozymes appear to have a distinct physiological distribution and function. IDO1 (EC 1.13.11.52), the focus of this study, is found in several tissues and is particularly over-produced in the placenta (*6–9*). The high placental abundance led to the initial discovery that IDO1 plays a pivotal role in immune tolerance, thereby protecting embryos from maternal immunity (*10,11*). Subsequent studies have revealed a role for IDO1 well beyond fetal development. Notably, IDO1 is produced at the highest levels in dendritic cells (DCs) (*12–16*) and to a lesser extent macrophages and eosinophils, where its expression is induced by lipopolysaccharide, interferon gamma, and other stimuli (*17-21*). The role of IDO1 in this immune cell population is generally believed to be twofold: *(i)* to starve invading microbes by depleting L-Trp and *(ii)* to produce metabolites initiating a signaling pathway that leads to immunosuppressive effects (*22*). The immunomodulatory properties are manifested in part through a suppression of the proliferation of cytotoxic and helper T cells and natural killer (NK) cells with concurrent upregulation of the production of regulatory T cells (Tregs) and myeloid-derived suppressor cells (*23-26*) (Figure 1). The extensive amount of data connecting IDO1 with a range of immunosuppressive outcomes, only briefly highlighted here, has led to the designation of IDO1 as a checkpoint modulator of immunity, both in terms of the immunogenicity and immune tolerance (*27-29*).

**Figure 1.**
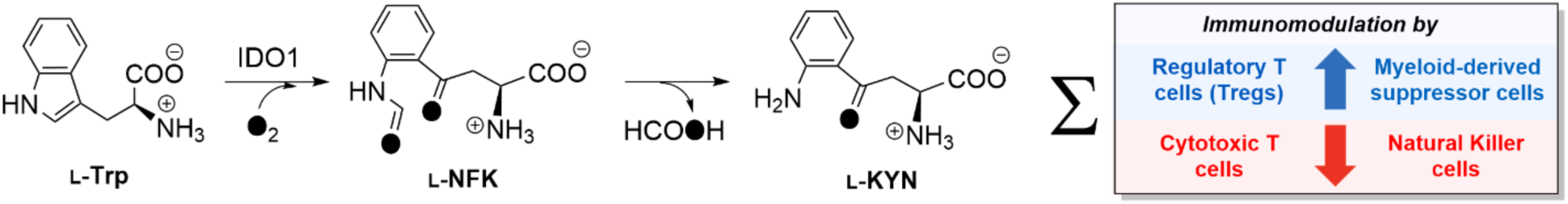
Reaction catalyzed by indoleamine 2,3-dioxygenase-1 (IDO1) and representative immunomodulatory outcomes linked to the IDO1. IDO1 catalyzes the L-tryptophan (L-Trp)-and O_2_-dependent formation of *N’*-formyl-L-kynurenine (L-NFK), which is hydrolyzed noncatalytically or by a distinct enzyme (arylformamidase, E.C. 3.5.1.9) to generate formic acid and L-kynurenine (L-KYN).

Initial studies with IDO1 isolated from the small intestine of *Oryctolagus cuniculus* (rabbit) led to the conclusion that the enzyme, initially called tryptophan pyrollase, is a hemoprotein that functions as a dioxygenase, i.e. incorporates both O atoms of O_2_ into the product L-NFK (*30-32*). Hemoproteins are a large family of mononuclear metalloproteins that contain an iron-bound porphyrin prosthetic group (i.e., heme) and can generally be categorized into two groups: those that directly engage with O_2_ to fulfill their physiological function or those that utilize H_2_O_2_ (*33*). The former can be further broken down into the O_2_-transport and storage proteins known as globins that includes myoglobin and hemoglobin and the O_2_-dependent enzyme catalysts that function as either oxygenases or oxidases. The functional diversity of hemoproteins is dictated primarily by variations in the residues that make up the heme active site environment, perhaps none more important than the axial iron ligand that resides on the proximal side of the porphyrin ring. This residue, which is critical for fine-tuning the reduction potential of the metal center, is typically Cys for O_2_-dependent enzymes that is exemplified by the well characterized family of cytochrome (CYP) P450 monooxygenases, which unlike IDO1, incorporate only one atom of O_2_ into a substrate while reducing the other to H_2_O (*34*). Studies with recombinant human protein (rhIDO1), first reported in 2000 (*35*), have identified a His as the proximal ligand (*36-38*). A proximal His is instead characteristic of the remaining types of hemoproteins: the H_2_O_2_-dependent peroxidases (with chloroperoxidase being a notable exception) and the O_2_-binding globin family (*33,39*). Thus, IDO1 appears unique among hemoproteins by not only functioning catalytically as a dioxygenase but also by exploiting a His proximal ligand for engaging O_2_ for catalysis.

Regardless of proximal ligand identity or the enzyme function, the commonly shared paradigm for O_2_-binding hemoproteins involves iron in the ferrous (+2) oxidation state prior to engagement of O_2_. For the globin family, the heme of the resting state—the default state before substrate binding— is predominantly the reduced, ferrous form. Contrastingly, the resting state of CYP P450 monooxygenases is predominantly in the oxidized, ferric heme. Consequently, CYP P450s require a separate reductase system to shuttle an electron into the heme to initiate substrate binding and the overall catalytic cycle. Evidence from in vivo and vitro studies suggest the resting state of human IDO1 is also the ferric heme (*35–38,40–42*). Based on the assumption that the established O_2_-binding dogma is followed by IDO1, a single-electron reduction to ferrous iron would be necessary for binding O_2_ and initiating catalysis. This would occur despite the lack of need for any electron input for completing a catalytic cycle, i.e. the overall IDO1-catalyzed reaction is redox neutral. In early studies with rabbit IDO1, reduction was achieved in vitro by the addition of two components (termed cofactors at the time of discovery): the synthetic dye methylene blue (MB) and ascorbic acid (AA) (*31,43*). Alternatively, ferric heme reduction could be bypassed by chemically or enzymatically generating superoxide (O_2_^•-^) (*43-45*). Reduced tetrahydrobiopterin (THBP) was also reported to substitute for MB/AA, thus providing a potential endogenous reductant for activating rabbit IDO1 for catalysis (*46*). Similarly to rabbit IDO1, human IDO1 uses the artificial MB/AA electron transport system to efficiently support catalysis (*47,48*). However, O_2_^•-^ does not appear to support human IDO1 activity (*49*). Additionally, and in contrast to the activity for many CYP P450 monooxygenases, rhIDO1 was shown to be inoperable using a peroxide shunt pathway that employs H_2_O_2_ and ferric enzyme as a substitute for O_2_ and the ferrous form of the enzyme (*50–52*). Indeed, assay conditions generally incorporate catalase to prevent H_2_O_2_-mediated inhibition. Polysulfides (*53*) and NADPH−cytochrome P450 reductase with or without cytochrome *b*_5_ (*49,54,55*) have also been reported to support rhIDO1 catalytic activity in vitro. However, whether these reduction systems operate in vivo to support IDO1 catalysis is still unclear.

Of the many striking features of the IDO1-catalyzed reaction, we were intrigued by the conclusions that IDO1 and homologs are the only members of the proximal His hemoproteins that use O_2_ as a substrate for a chemical transformation within the biological milieu. This was particularly interesting given the inability to definitively identify a palatable, endogenous “activator” (aka initiating factor) responsible for ferric heme reduction and hence O_2_ engagement. We therefore set out to find operational parameters that, of critical importance, adequately reflect the physiological function of IDO1. Here we report the identification of a catalytically competent peroxidase-like shunt pathway for IDO1 that satisfies the activity-function connection. The key finding as reported herein is that sodium hypochlorite (NaOCl), aka bleach, activates ferric IDO1 in a catalytic manner. Further biochemical studies revealed that O_2_ remains essential when using hypochlorite and that other hypohalites can support catalysis. The driving rationale and the implications for the discovery of hypochlorite-mediated and O_2_-dependent catalysis from the perspective of IDO1’s physiological function in immunosuppression is discussed. Furthermore, a catalytic mechanism, which is not only consistent with the data presented here but is backed by strong literature precedent, is proposed that represents a new modus operandi for O_2_ activation for hemoproteins.

## RESULTS

### Hypochlorite supports rhIDO1 catalysis

Activity of recombinant human IDO1 (rhIDO1) was initially examined in reactions using the artificial MB/AA electron transport system followed by heat workup to remove protein (Figure 2a). As expected, a new peak appeared in the HPLC-MS chromatogram that coincided with the disappearance of the L-Trp peak (Figure 2b). HRMS of the new peak, which had a distinct UV/VIS spectrum than reported for the known product of IDO1 (L-NFK), was consistent with the formation of (*E*)-4-[2-(formylamino)phenyl]-4-oxo-2-butenoic acid, the deamino derivative of L-NFK (dNFK) (Figure 2c). Sample work-up using ultrafiltration or trichloroacetic acid-mediated protein precipitation instead of boiling yielded a predominant peak eluting before L-Trp with UV/VIS and mass spectra consistent with L-NFK, demonstrating that dNFK is a degradation product due to heat treatment under mildly alkaline conditions. For reproducibility, all subsequent reactions were terminated by boiling, forming predominantly dNFK, but product yield was calculated based on both products (dNKF + unconverted L-NFK). By analyzing reactions following the complete conversion of several concentrations of L-Trp to dNFK + L-NFK [χ_321nm_ = 3750 M^-1^cm^-1^ (56)], the extinction coefficient of dNFK at 280 nm was determined to be 3075 M^-1^cm^-1^.

**Figure 2.**
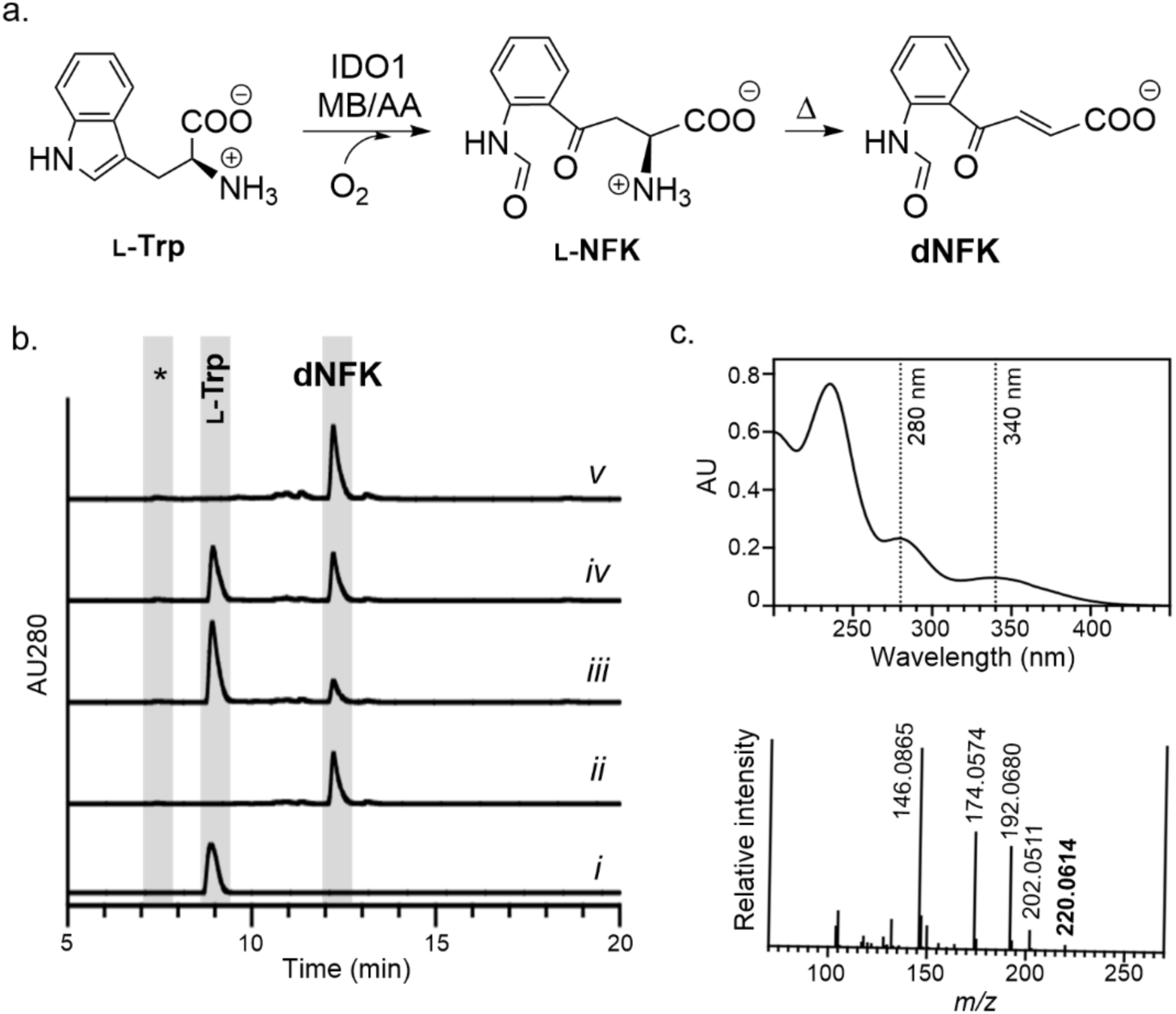
Reaction catalyzed by indoleamine 2,3-dioxygenase-1 (IDO1). (a) The established IDO1 reaction using methylene blue (MB) and ascorbic acid (AA) to form *N’*-formyl-L-kynurenine (L-NFK), which can degrade to the deamino derivative dNFK. (b) HPLC traces of the rhIDO1-catalyzed reaction including the control (no enzyme, trace *i*); rhIDO1 reaction for 2.5 h using established conditions with MB/AA (trace *ii*); rhIDO1 reaction for 2.5 h with 1 mM L-Trp, 100 µM NaOCl, and 10 mM AA (trace *iii*); rhIDO1 reaction for 2.5 h with 1 mM L-Trp, 100 µM Ca(OCl)_2_, and 10 mM AA (trace *iv*); and rhIDO1 reaction for 14 h with 1 mM L-Trp, 100 µM Ca(OCl)_2_, and 10 mM AA (trace *v*);. AU280, absorbance units at 280 nm; *, retention time for L-NFK, which has minimal absorbance at 280 nm. (c) Spectroscopic analysis of dNFK including the UV/VIS spectrum and high-resolution mass spectrum in positive mode (calculated *m/z* = 220.0610 for [C_11_H_9_NO_4_ + H]^+^ ion corresponding to dNFK).

Neutrophils and macrophages undergo a respiratory burst in response to infection. This burst pathway involves three enzymes, NADPH oxidase, superoxide dismutase, and myeloperoxidase, that sequentially convert O_2_ to O_2_^•-^, H_2_O_2_, and HOCl, all of which are considered reactive oxygen species (ROS) capable of damaging invading microbes. Although often underappreciated, HOCl has also been demonstrated to have immunomodulatory properties via an unknown mechanism that alters cytokine signaling and suppresses inflammation in a manner that is uncannily like IDO1. With this insight and the knowledge that other heme-dependent enzymes (but, as previously noted, not rhIDO1) are known to use O_2_^•-^ or H_2_O_2_ to bypass the need for ferric heme reduction, we tested the potential for ferric-heme rhIDO1 to use HOCl to catalyze the formation of L-NFK from L-Trp. HPLC analysis of rhIDO1 reactions using sodium hypochlorite (NaOCl; 100 µM) revealed reproducible conversion of L-Trp (1 mM) to dNFK. Freshly prepared hypochlorite solution from solid Ca(OCl)_2_ was able to substitute for NaOCl. To confirm the reaction, lactoperoxidase (LPO; EC 1.11.1.7); an enzyme found in mammalian secretions such as milk, saliva, tears, and mucus which is commercially available from bovine milk; was employed for in situ generation of HOCl. Incubations of rhIDO1 and LPO with H_2_O_2_ and excess NaCl revealed formation of dNFK, although yields were substantially less in comparison to reactions starting directly with HOCl. Consistent with prior reports, rhIDO1 was inactive when substituting HOCl with H_2_O_2_ (Table S1) or using xanthine oxidase/xanthine to generate O_2_^•-^ (and potentially ^•^OH) in situ (57). In total, the data demonstrates that HOCl can support oxygenation for a heme-dependent enzyme analogous to the peroxide shunt reaction, although for rhIDO1 the reaction is highly specific for HOCl when compared to other endogenous ROS species.

### Characterization of rhIDO1 activity with HOCl and potential substitutes

rhIDO1 reactions with 100 µM NaOCl and variable [AA] revealed a steady decline in enzyme activity when [AA] < 8 mM (Figure 3a), which was previously observed for IDO1 when using MB. At higher concentration of NaOCl (1 mM), substantially less AA was necessary for complete conversion to product (Figure 3a). Notably, at both concentrations of hypochlorite, reactions proceeded even in the absence of AA, suggesting AA enhances but is not essential for activity (Figure 3b). At constant AA (5 or 10 mM), an increase in activity was likewise observed upon increasing the concentrations of NaOCl or Ca(OCl)_2_ with comparable conversion on a ^-^OCl equivalency basis (Figure 3c and d). In contrast to reactions with variable [AA], however, no product was detected in the absence of HOCl. Furthermore, increasing the concentration of HOCl above 1 mM produced HPLC spectra with decreased substrate/product intensities and increased noise, probably due to the non-specific oxidative degradation of L-Trp and/or products. Using optimal concentrations, the highest product conversion was observed at pH of 7.4 (Figure S1). Notably, the equilibrium ^-^OCl +H_2_O⇋HOCl + ^-^OH (p*K*a = 7.5) exists roughly as a 1:1 mixture of ^-^OCl and HOCl at pH = 7.4. For simplicity, all chlorine species are referred to as HOCl.

**Figure 3.**
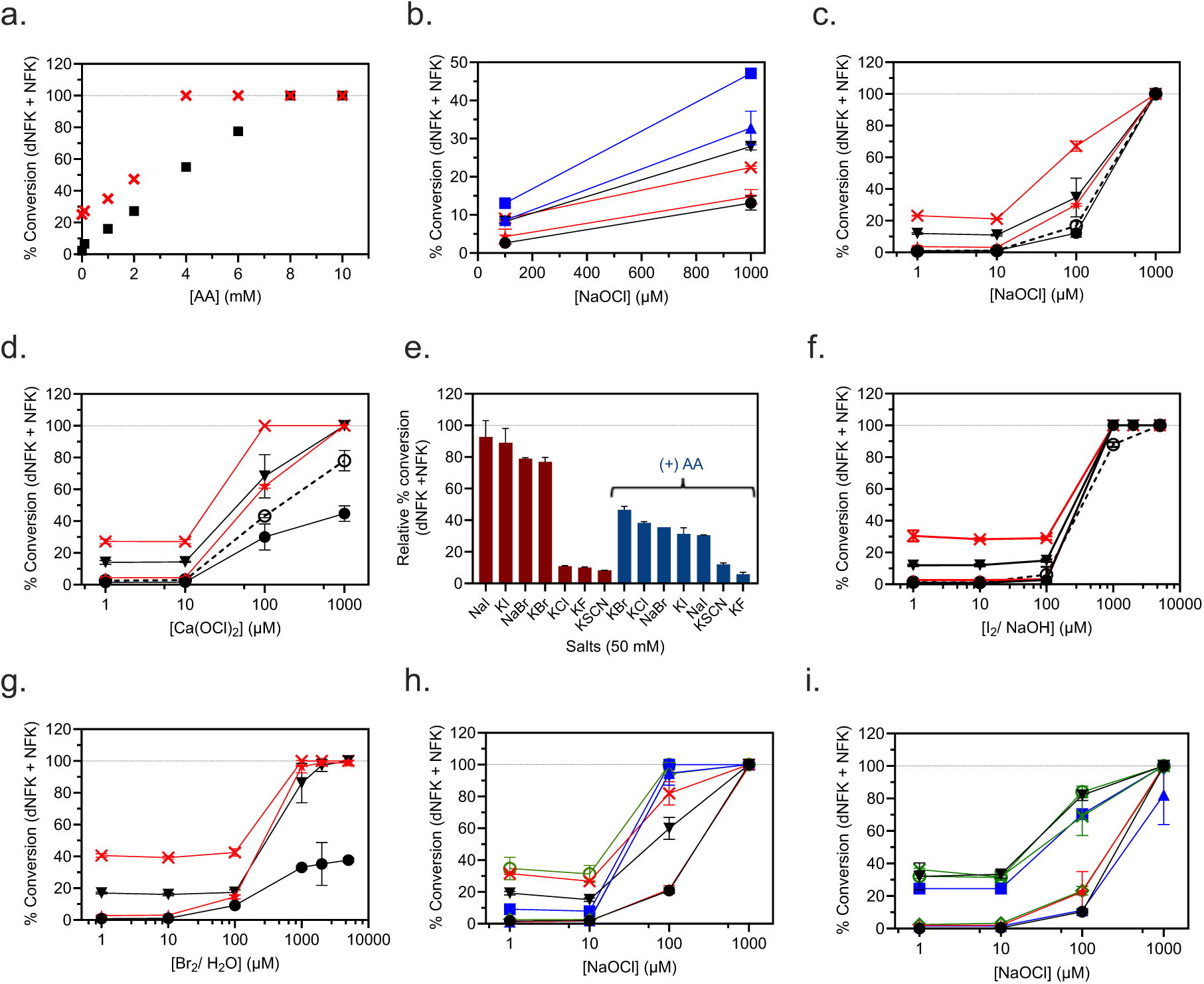
Characterization of recombinant human indoleamine 2,3-dioxygenase-1 (rhIDO1) with (pseudo)hypohalites. (a) Reactions with variable [ascorbic acid (AA)] and [NaOCl] = 100 mM ( ▪ ) or 1 mM ( 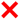 ). The dashed line represents 100% conversion of 1 mM L-Trp to products. (b) Reactions in the absence of AA and variable [NaOCl] without additional salts (**black** data points) after 2.5 h ( ● ) and 18 h ( ▾ ); with 1 mM NaI (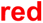 data points) after 2.5 h ( ★ ) and 18 h ( 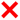 ); or with 10 mM NaI (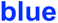 data points) after 2.5 h ( 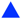 ) and 18 h ( 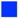 ). Note that trend lines are depicted and the y-axis scale is less than 100% conversion in order to better visualize the difference between the data points. (c) Semi-logarithmic plot for reactions with variable [NaOCl] with 5 mM AA after 2.5 h ( ● ) and 18 h ( ▾ ) or with 10 mM AA after 2.5 h ( ★ ) and 18 h ( 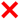 ); or reactions with 10 mM AA after 2.5 h ( ○ ) with rhIDO1 pre-treated with ferricyanide (dashed line). (d) Semi-logarithmic plot for reactions with variable [Ca(OCl)_2_] with 5 mM AA after 2.5 h ( ● ) and 18 h ( ▾ ) or with 10 mM AA after 2.5 h ( ★ ) and 18 h ( 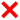 ); or reactions with 10 mM AA after 2.5 h ( ○ ) with rhIDO1 pre-treated with ferricyanide (dashed line). (e) Reactions using lactoperoxidase, H_2_O_2_, and indicated salt for the in situ generation of (pseudo)hypohalites, in the absence or presence of AA. (f) Semi-logarithmic plot for reactions with variable [I_2_/ NaOH] with 5 mM AA after 2.5 h ( ● ) and 18 h ( ▾ ) or with 10 mM AA after 2.5 h ( ★ ) and 18 h ( 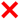 ); or reactions with 10 mM AA after 2.5 h ( ○ ) with rhIDO1 pre-treated with ferricyanide (dashed line). (g) Semi-logarithmic plot for reactions with variable [Br_2_ (*aq*)] with 5 mM AA after 2.5 h ( ● ) and 18 h ( ▾ ) or with 10 mM AA after 2.5 h ( ★ ) and 18 h ( 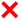 ). (h) Semi-logarithmic plot for reactions with variable [NaOCl] with 10 µM NaI after 2.5 h ( ● ) and 18 h ( ▾ ); with 100 µM NaI after 2.5 h ( ★ ) and 18 h ( 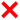 ); with 1 mM NaI after 2.5 h ( 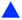 ) and 18 h ( 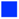 ); or with 10 mM NaI after 2.5 h ( **◇** ) and 18 h ( ○ ). (h) Semi-logarithmic plot for reactions with variable [NaOCl] with 10 µM NaBr after 2.5 h ( ● ) and 18 h ( ▾ ); with 100 µM NaBr after 2.5 h ( ★ ) and 18 h ( 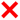 ); with 1 mM NaBr after 2.5 h ( 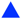 ) and 18 h ( 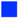 ); or with 10 mM NaBr after 2.5 h ( **◇** ) and 18 h ( ○ ).

To probe halide selectivity, we next examined rhIDO1 activity using an LPO-coupled hypohalite/ hypothiocyanite generation system. LPO is known to catalyze the oxidation of Cl^-^, Br^-^, I^-^, or the pseudohalide SCN^-^ in the presence of H_2_O_2_, thereby producing hypohalites or hypothiocyanite, respectively. All four (pseudo)halides successfully supported IDO1 catalysis, with the order of activity as follows: NaI ≈ KI > NaBr ≈ KBr >>> KCl ≈ KF ≈ KSCN (Figure 3e). This order is identical to the reported selectivity (based on second order rate constants) for bovine LPO alone except for one notable difference: KSCN is the preferred substrate for LPO (58). Thus, rhIDO1 appears selective toward true hypohalites. In contrast to reactions solely with rhIDO1, better conversion was observed in the absence of AA, suggesting AA potentially inhibits LPO, which has not been previously reported. Furthermore, the rhIDO1 reaction starting with NaOCl was inhibited by H_2_O_2_ as previously reported using MB/AA assay conditions. To avoid potential interference due to incompatibility with the LPO-coupled reaction conditions, rhIDO1 activity was tested with bromine water [Br_2_ (*aq*)] and fresh preparations of I_2_ (*s*) in water as a source of hypobromite and hypoiodite, respectively. Both solutions generate hypohalites that are formed via several pH-dependent reactions that are driven by kinetic and thermodynamic parameters. As expected, aqueous Br_2_ and I_2_ were effective substitutes for the LPO-coupled system with catalytic behavior identical to HOCl. However, the reaction rate and overall progress with 1 mM I_2_ in solution was significantly greater than those observed with HOCl (Figure 3f). Assuming the equilibrium I_2_ + H_2_O ⇋ HOI + I^-^ + H^+^ with *K_eq_* = 5.4 x 10^-13^ M^2^ dominates, reactions with1 mM I_2_(aq) at pH 7.4 are equivalent to an initial concentration of [HOI]_i_ = ∼110 µM (the p*K*_a_ for HOI is estimated to be between 10.5-11). This amount of HOI in the presence of 5 mM AA and 1 mM L-Trp resulted in complete conversion of L-Trp to dNFK after 2.5 h (Figure 3f). Comparatively, conversion with 100 μM NaOCl was 12% (Figure 3c). With bromine water, assuming Br_2_ + H_2_O ⇋ HOBr + Br^-^ + H^+^ with *K_eq_* = ∼5 x 10^-9^ M^2^ so that [HOBr]_i_ = ∼992 µM (the p*K*_a_ for HOBr is 8.65), 2.5 h reactions with 5 mM AA gave 30% conversion which was increased to 100% with 10 mM AA (Figure 3g). Comparatively, conversion with 1 mM NaOCl was 100% at both AA concentrations. As noted above with NaOCl, reactions with concentrations > 1 mM of hypohalite source were difficult to interpret due to the loss of UV active species in the HPLC chromatograms. Finally, HOI and HOBr were generated in situ by reacting NaOCl with NaI and NaBr, respectively. Nearly complete conversion was realized within 2.5 h when starting with 100 μM NaOCl and 1 mM L-Trp in the presence of 1 mM NaI (Figure 3h). Comparatively, conversion with 1 mM NaBr was 12% (Figure 3i), which is identical to reactions in the absence of NaBr (only NaOCl). Additionally, inclusion of NaI with NaOCl in reactions without AA approximately doubled the conversion of L-Trp to product (Figure 3b).

Taking into consideration the apparent distinct functional characteristics between IDO1 isolated from rabbit intestine versus the recombinant human form, the recombinant version from *Canis lupus dingo* was produced and purified from *E. coli*. A preliminary assessment revealed *Cd*IDO1, identically to rhIDO1, catalyzed the conversion of L-Trp to L-NFK using representative hypohalites (Figure S2). Thus, hypohalite-dependent IDO1 activity extends beyond humans and is potentially conserved across mammals.

### Kinetic and TTN analysis

Single-substrate kinetic analysis with respect to variable [L-Trp] was next examined with the cofactors/activators that support rhIDO1 catalysis. The velocity plots followed Michaelis-Menten kinetics regardless of condition (Figure 4); the extracted kinetic parameters are summarized in Table 1. The second order rate constants are relatively close in value regardless of the identity or concentration of the cofactor/activator. An increase in *K_m_* with respect to L-Trp is observed with increasing concentration of HOCl, which was off-set by a comparable fold-increase in *k_cat_*. Consistent with our end-point assays suggesting no H_2_O_2_ inhibition, catalase—which catalyzes the conversion of H_2_O_2_ to H_2_O and O_2_ and is generally included in IDO1 assays by others—had no effect on the kinetic parameters when rhIDO1 was assayed with NaOCl. Finally, and likewise consistent with reported kinetic assays employing MB/AA, rhIDO1 catalyzes the reaction with D-Trp but has a significantly higher *K_m_* when compared to the proteinogenic amino acid L-Trp.

**Figure 4.**
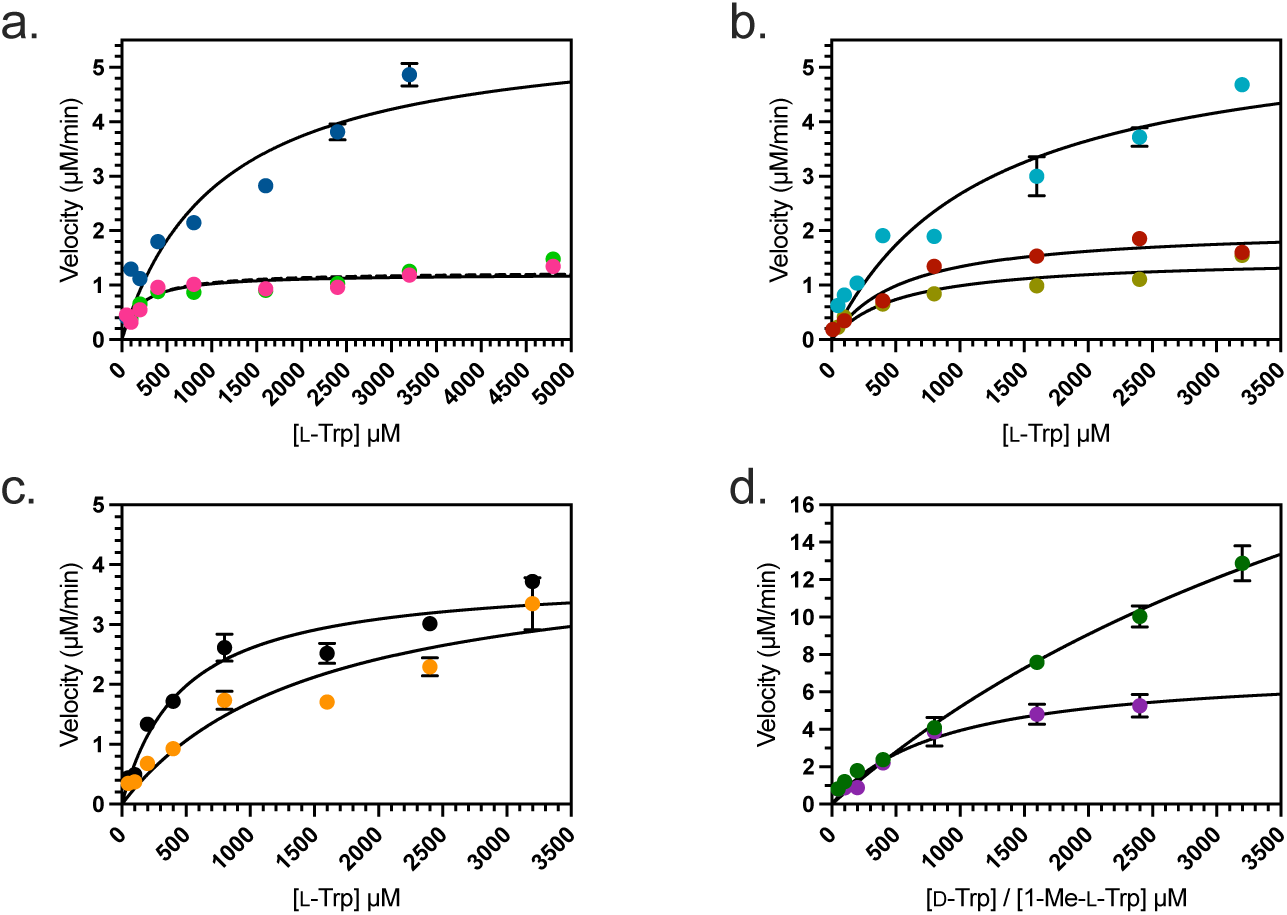
Michaelis-Menten plots for a single-substrate kinetic analysis of rhIDO1. rhIDO1 reactions with variable [L-Trp] and (a) 200 µM NaOCl ( 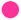 ), 1 mM NaOCl ( 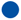 ), and 200 µM NaOCl with 10 µg/mL catalase ( 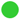 ); (b) 100 µM Ca(OCl)_2_ ( 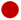 ), 500 µM Ca(OCl)_2_ ( 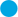 ), and 100 µM Ca(OCl)_2_ and 100 µM NaBr ( 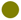 ); (c) 1 mM I_2_/ NaOH ( 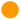 ), 1 mM Br_2_/ H_2_O ( **●** ); (d) with variable [D-Trp] and 100 µM Ca(OCl)_2_ ( 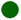 ), and with variable [1-Me-L-Trp] and 200 µM NaOCl ( 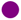 ).

**Table 1.** Single substrate kinetic parameters for rhIDO1.

| Substrate | Activator/Cofactor<br>(concentration) | $K_m$ (M) | $k_{cat}$ (min<sup>-1</sup>) | $k_{cat}/K_m$<br>(relative) |
| --- | --- | --- | --- | --- |
| L-Trp | NaOCl (200 μM) | $(1.8 \pm 0.5) \times 10^{-4}$ | $6.0 \pm 0.4$ | 1 |
| | NaOCl (200 μM) + catalase<sup>a</sup> | $(2.0 \pm 0.6) \times 10^{-4}$ | $6.2 \pm 0.4$ | 0.94 |
| | Ca(OCl)<sub>2</sub> (100 μM) | $(5.5 \pm 1.2) \times 10^{-4}$ | $10.4 \pm 0.7$ | 0.56 |
| | NaOCl (1 mM) | $(1.1 \pm 0.4) \times 10^{-3}$ | $29 \pm 4$ | 0.79 |
| | Ca(OCl)<sub>2</sub> (500 μM) | $(1.2 \pm 0.4) \times 10^{-3}$ | $29 \pm 4$ | 0.73 |
| | I<sub>2</sub> (1 mM) | $(1.5 \pm 0.6) \times 10^{-3}$ | $21 \pm 4$ | 0.42 |
| | Br<sub>2</sub> (1 mM) | $(4.7 \pm 0.9) \times 10^{-4}$ | $19 \pm 1$ | 1.2 |
| | Ca(OCl)<sub>2</sub> (100 μM) + NaBr<sup>b</sup> | $(5.2 \pm 1.8) \times 10^{-4}$ | $8.0 \pm 0.7$ | 0.46 |
| D-Trp | Ca(OCl)<sub>2</sub> (100 μM) | $\geq (5.9 \pm 1.6) \times 10^{-3}$ | $\geq 180 \pm 35$ | 0.9 |
| 1-Me-L-Trp | NaOCl (200 μM) | $(8.6 \pm 2.1) \times 10^{-4}$ | $37 \pm 4$ | 1.3 |

<sup>a</sup>10 μg/mL; <sup>b</sup>100 μM.

### Reassessment of 1-methyl-L-tryptophan, a reported IDO1 inhibitor and substrate

The substrate analogue 1-Me-DL-Trp has been described as a competitive inhibitor and alternative—albeit poor—substrate with respect to L-Trp when assayed with rhIDO1 and the MB/AA conditions (59–61). Using the newly identified reaction conditions (200 µM NaOCl and heat workup), rhIDO1 reactions with commercially available 1-Me-L-Trp resulted in the loss of the peak corresponding to 1-Me-L-Trp and the appearance of a distinct peak with a λ_max_ at 227 nm and a shoulder at 278 nm with a broad tail extending to longer wavelengths (Figure S3). HRMS and NMR analyses of the purified product were consistent with the identity of the new peak as (*E*)-4-[2-(*N*-methylformamido)phenyl]-4-oxo-2-butenoic acid (*N*-Me-dNFK) (Figure S4). Inspection of the NMR spectrum suggested the product existing as a mixture of rotamers (2:1):(*E*:*Z*) at room temperature due to the restricted rotation about the N–C(O) bond. Increasing the temperature during acquisition of the ^1^H-NMR spectrum resulted in the convergence of peaks at ∼6.60 and ∼7.35 ppm, providing evidence for this mixture of rotamers (Figure S4g). Furthermore, the *cis* relationship of the *N*-methyl group and the formyl hydrogen of the minor conformer was supported by NOESY (Figure S4f). Single-substrate kinetic analysis using HPLC to detect the product (Figure 4d and Table 1) revealed *K_m_* = (8.6 ± 2.1) x 10^-4^ M^-1^ and *k_cat_* = 37 ± 4 min^-1^. This equates to a slight increase (1.3-fold) in the second order rate constant when compared to L-Trp under identical reaction conditions.

The detection of the putative rhIDO1 product prior to heat-induced deamination, *N’*-Me-L-NFK, was attempted using two prior methods routinely employed to quantitate rhIDO1 activity. Using the identical protocol with Ehlrich’s reagent as described (59), the absorbance of the resulting chromophore at 480 nm was 9-times less than expected despite the completion of the reaction as judged by co-analysis using HPLC. Similarly, the extinction coefficient at 321 nm for *N’*-Me-L-NFK was calculated to be 408 M^-1^cm^-1^ (Figure S3), ∼9-times less than the extinction coefficient for L-NFK (χ_321 nm_ = 3750 M^-1^cm^-1^) that we assume was previously used to calculate *N’*-Me-L-NFK concentration (61). In total, the data is consistent with past experiments underestimating the activity of IDO1 with *N’*-Me-L-NFK.

### Preliminary mechanistic insight

As previously noted, complete conversion of 1 mM L-Trp to product was achieved with rhIDO1 (1-2 mM) and sub-stoichiometric (<1 mM) amounts of HOCl. By lowering the rhIDO1 concentration (0.05 mM), the total turnover number (TTN) was estimated as ≥ 2000 when initiating the reaction with 100 µM or 500 µM of Ca(OCl)_2_. Overall, this data suggests HOCl is not a substrate and is instead consistent with a role as an activator, i.e. gets consumed during the first catalytic cycle but is not required in subsequent cycles, or a cofactor, i.e. gets regenerated while participating in every catalytic cycle. This subsequently suggests that, as previously reported by others using MB/AA assay conditions, the source of both O atoms remains O_2_ without incorporation of the O atom of the hypohalite. We attempted several times to demonstrate this using isotopic enrichment experiments following literature protocol and variations thereof but quickly realized that the solvent exchange with L-NFK was too rapid to provide meaningful data in our hands. Consequently, we simply examined whether O_2_ was indeed required for reactions using the hypohalite conditions. For this purpose, solid I_2_ was added to a thoroughly deoxygenated solution containing L-Trp, AA, and rhIDO1 and allowed to react under a nitrogen atmosphere for 4 hr. LCMS analysis revealed 7% of the starting material was converted to the product (Figure S5). In contrast, complete substrate conversion occurred in the control reaction, wherein the deoxygenated mixture (including IDO1) was opened to an O_2_ atmosphere after I_2_ addition (Figure S5). These results indicate that O_2_ is still required for rhIDO1 catalysis in hypohalite conditions and likely remains the source of both O atoms in L-NFK, hence functioning as a dioxygenase.

Finally, rhIDO1 retained ≥80% of its activity following treatment with 1 mM potassium ferricyanide (III) (Figures 3b, c, and f – dashed lines). Therefore, HOCl-dependent rhIDO1 activity is unlikely to be initiated in the ferrous form despite still requiring O_2_ to catalyze the conversion of L-Trp to L-NFK.

## DISCUSSION

IDO1 catalyzes the O_2_- and heme-dependent conversion of L-Trp to L-NFK to initiate the kynurenine pathway. Previous studies with IDO1 have independently revealed at least three different electron reduction systems that are able to initiate IDO1 catalysis in vitro, namely MB/AA (48), hydrogen sulfide and polysulfides (53), or NADPH-cytochrome P450 reductase with or without cytochrome *b_5_* (49, 55). The function of each of these systems has been ascribed to regenerating and maintaining ferrous IDO1, which has been deemed the necessary form to achieve the dioxygenase activity. The data provided here demonstrate yet another way to achieve the rhIDO1-catalyzed conversion of L-Trp to L-NFK: catalytic initiation with hypohalous acids. This hypohalous acid-dependent activity is distinct from the other activation systems in that it altogether bypasses the need for generating the ferrous rhIDO1. Instead, this activation is reminiscent of peroxidases or the CYP P450 peroxide shunt pathway, although with key proposed mechanistic differences that are highlighted below. Nonetheless, to our knowledge, this is the first demonstration of a human enzyme that employs hypohalous acids for catalytic activation.

A major driving factor for examining rhIDO1 activity with hypohalous acids was consideration of the biological milieu that rhIDO1 is expected to function. A particularly formative driver was the work by Odobasic *et al*. that revealed MPO and more specifically, its product hypochlorous acid (and, importantly, not hydrogen peroxide) attenuates T cell driven tissue inflammation by “suppressing” the function of dendritic cells in draining lymph nodes (62). One of the many important findings of this work included that human monocyte-derived dendric cells cultured with neutrophil supernatants or MPO led to a significant decrease in HLA-DR and CD86 expression, both markers for productive T cell priming. Although many gaps remain to be addressed, the study provided compelling support for HOCl-driven immunomodulation via dendritic cell function. Likewise, studies have shown topical HOCl reduces inflammation and the pro-inflammatory response for many skin conditions including eczema, suggesting a role for rhIDO1 in this milieu as well (63). By activating rhIDO1, our data further supports that endogenous HOCl is more than a microbicidal agent, which is the predominant functional significance put forth in the community. In addition to these observations of HOCl immunomodulation, another driving factor for testing HOCl was the consideration of the three “flavors” of endogenous reactive oxygen species that are enzymatically formed during an immune response: O_2_^•-^, H_2_O_2_, and HOCl (64). Interestingly, HOCl is often omitted and replaced with ^•^OH as one of these three primary ROS (64,65). Both O_2_^•-^ and H_2_O_2_ (via the peroxide shunt for CYP P450s or naturally with peroxidases) are known to substitute for O_2_ and bypass the requisite 1-*e*^-^reduction for initiating catalysis for many heme-containing enzymes (66). Although previously tested for binding and partial reactivity with myoglobin, horseradish peroxidase, and bovine catalase (67) and demonstrated as a reactive substrate with synthetic porphoryin mimics (68), HOCl is rarely (if ever) discussed as a potential oxidant for initiating heme-dependent chemistry. However, here we have shown that HOCl is the only ROS “flavor” used by rhIDO1 to generate L-NFK. Thus, it is tempting to speculate, at least in the scenario for dendritic cell function in draining lymph nodes, that HOCl functions as a signaling molecule by first entering dendritic cells, the established predominant factory for rhIDO1 production, and then activating rhIDO1 for concomitant depletion of L-Trp and formation of L-NFK. The presence or absence of these metabolites in turn ultimately regulate the differentiation of naïve T cells, a well-established connection to rhIDO1 activity.

The discovery that rhIDO1 uses HOCl to initiate multiple catalytic cycles re-opens the door for a new round of mechanistic studies on this intriguing enzyme family and potentially other hemoproteins. While we initially considered the possibility that the two O atoms of L-NFK originate from two HOCl equivalents for each catalytic cycle, the data are consistent with sub-stoichiometric HOCl with respect to L-Trp, suggesting HOCl is either a cofactor (i.e., re-formed at the end of each catalytic cycle) or activator. Furthermore, the HOCl-initiated reaction still requires O_2_. Unfortunately, in contrast to reports by others (32,69), rapid solvent exchange of L-NFK prevented isotopically tracking the O atoms with acceptable rigor in our hands. Consequently, at this stage, we propose that HOCl initiates the catalytic cycle by ligation to the ferric heme in the distal pocket, which leads to a two-electron transfer to form Cl^-^ and a ferryl-oxo porphyrin radical cation, Compound I. From Compound I, minimally two pathways can be envisioned. The first exploits the oxidizing power of Compound I for abstraction of a π-electron from substrate L-Trp leading to a carbon-centered radical cation that directly reacts with triplet O_2_. The subsequent peroxyl L-Trp radical is reduced followed by dioxetane formation leading to the rearrangement to form L-NFK. Reactions between carbon-centered radicals and O_2_ to form hydroperoxides are known to occur at rates near diffusion (*k* = 10^9^-10^10^ M^-1^s^-1^) in solution (70). The reaction of O_2_ during H_2_O_2_-mediated catalysis using metmyoglobin and methemoglobin from several sources lends support for this chemistry, as EPR studies suggest the formation of peroxyl radical at C3 of a L-Trp residue, which is likewise consistent with the formation of a C3 radical of the indole for rhIDO1 following HOCl-dependent Compound I genesis (71–74). Not surprisingly, the major peroxidation product of this L-Trp residue has been shown to be L-NFK (75). The intermediacy of a hydroperoxy species is further supported by analogous results examining chemical-sensitized photooxygenation (via the more relevant type I free radial mechanism or, alternatively, a type II singlet O_2_ mechanism) of indole derivatives (76–78) including L-Trp residues in non-heme proteins (70, 79).

A provocative, alternative pathway for O_2_ engagement is the direct reaction of the ferryl-oxo of Compound I with O_2_ to transiently generate ozone (O_3_) (Figure 5, path b). In this mechanism the O atom of the ferryl oxo could potentially be first reduced to the oxide radical (•O^-^) that then reacts with triplet O_2_ to form a heme-ligated transient ozonide radical anion (•O_3_^-^) and Compound II (not shown), the former of which is rapidly oxidized to O_3_ with concomitant reduction of the latter to ferric rhIDO1. Importantly for the initiation of path b, experimental and computational analysis of horseradish peroxidase are consistent with a Compound I ground state having a triplet spin pair on the ferryl-oxo unit (80–83), suggesting that such a radical coupling is minimally spin-allowed. Subsequent ozonolysis of L-Trp to L-NFK would follow, a reaction which mimics the chemical synthesis employed for making L-NFK from L-Trp (84) and was determined chemically feasible in aqueous, neutral conditions by us (Figure S6). The IDO1-catalyzed reaction could potentially proceed through a molozonide and peroxide intermediates as shown, which is based on NMR analysis of the synthetic reactions. Peroxide reduction by Cl^-^ to form L-NFK would regenerate the HOCl-ligated ferric rhIDO1. Central to this mechanism is the ability of rhIDO1 to generate O_3_, an entity that is typically considered in the context of atmospheric chemistry but not biological systems. Although the standard reduction potential of O_3_ is indeed relatively high, it is still less than the hydroxyl radical, •OH (notably of which is the conjugate acid of the oxide radical that is proposed here) that has been reported as a product of several hemoproteins when exposed to H_2_O_2_ under suboptimal or uncoupling conditions, including horseradish peroxidase (85) and others. The estimated difference in standard reduction potential in water between H_2_O_2_ and •OH (1.78 V and 2.80 V, respectively) is significantly greater than HOCl and O_3_ (1.48 V, acid form and 2.08 V, respectively), suggesting the HOCl-O_3_ couple is feasible when simply considering the comparable thermodynamic barriers (86). Finally, while ozonolysis has been demonstrated to convert L-Trp residues to the corresponding L-NFK residues and degrade heme prosthetic groups in vitro (87–89), the endogenous production of O_3_, and the potential for an in vivo function, has not been widely accepted. Several noncatalytic processes using alternative ROS have been proposed to generate O_3_ (90), but perhaps most intriguing is the proposal of catalytic antibody-mediated O_3_ production (91–93) and that the HOCl-producing enzyme MPO plays an important role in this catalytic process (93). With respect to function, this ozone generation has been proposed to occur in neutrophils, consistent with a role in the immune response. In support of an immunosuppressive role for O_3_, it is interesting to note that medical ozone therapy, although not FDA approved in the US, has been used throughout the world to manage many chronic inflammatory diseases including MS and arthritis.

**Figure 5:**
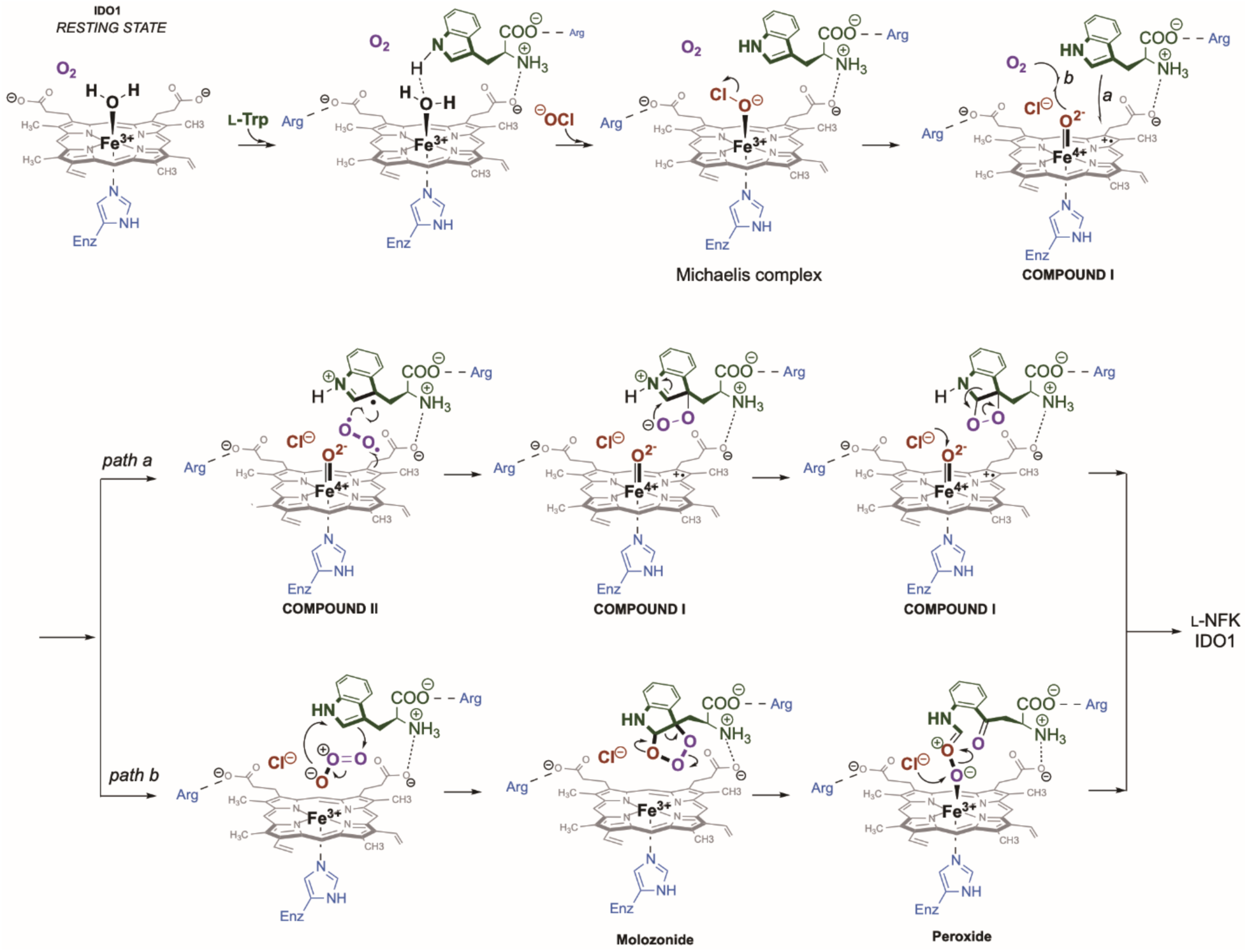
Hypothetical mechanism for rhIDO1-catalyzed conversion of L-Trp to L-NFK, with NaOCl as a catalytic cofactor/activator. The proposed binding interactions are based on several independent structures of rhIDO1, although protein dynamics likely alter these interactions during the catalytic cycle, which in turn is predicted to modulate the reducing power of the heme prosthetic group. The formation of Compound I mimics the peroxide shunt pathway that is realized with several hemoproteins. Compound I is proposed to proceed either through path a, forming a dioxetane intermediate or through path b, following an ozonolysis mechanism. In both routes, chloride is shown as the final reducing agent that regenerates heme-bound hypochlorite, thus behaving as a true cofactor. However, the possibility that the next cycle starts without regeneration of HOCl, hence behaving as a catalytic activator.

Following the discovery of the HOCl-dependent IDO1 activity, we initiated studies to re-assess perceived substrates and inhibitors that were identified using the MB/AA assay system. Here we report the results from one of these, 1-Me-DL-Trp, which has been the source of confusing if not contradicting data during its development as therapeutic agent. When human rIDO1 was first characterized in vitro, 1-Me-DL-Trp was concluded to be a poor substrate relative to L-Trp (7% relative conversion to an unidentified product) and a relatively strong inhibitor (limited L-Trp conversion to 27%) when using Ehlrich’s reagent to detect product formation (59). A later study using the same workup conditions revealed differences between the enantiomers, with 1-Me-L-Trp behaving as a competitive inhibitor with respect to L-Trp (*K_i_* = 19 mM) unlike the D-isomer (*K_i_* > 100 mM) (60). Unexpectedly, the L-isomer, in contrast to the D-isomer, was shown to have poor immunosuppression properties in vivo in the same study (60).

A more recent assessment of rhIDO1 led to the conclusion that 1-Me-L-Trp is indeed a substrate with the identity of the product being supported, for the first time, by low resolution LC-MS (87). Single-substrate kinetic analysis by monitoring product formation at 321 nm suggested that 1-Me-L-Trp was a relatively poor substrate with *K_m_* = 150 +/-11 µM and *k_cat_* = 0.027 +/- 0.001 s^-1^, approximately 9 x 10^-4^ the catalytic efficiency in comparison to the reaction with L-Trp (61) Using HPLC with diode array as a primary detection method along with a thorough spectroscopic analysis of the degradation product, we confirmed here that 1-Me-L-Trp is indeed a substrate using the newly discovered HOCl-dependent assay conditions. However, our data suggests 1-Me-L-Trp is actually as good as, if not better substrate than L-Trp. With this realization, the detection methods previously employed were reassessed, which revealed that *N’*-Me-L-NFK formation was likely greatly underestimated. Based on these results, it is thus tempting to speculate that prior failure to move 1-Me-L-Trp forward as an immunomodulatory therapeutic is a consequence of being efficiently converting 1-Me-L-Trp in vivo by rhIDO1, potentially to a product that behaves similarly in vivo to L-NFK. It will now be interesting to test whether this is indeed the case. Furthermore, we expect that the assay blueprint established here can now serve as the foundation for re-examining other categorized inhibitors and/or substrates, including D-isomer of 1-Me-Trp. Finally, from a mechanistic perspective, the turnover improvement with 1-Me-L-Trp is speculated to be attributed to the electron-donating group inductive effect, which supports the reactivity of hypothetical intermediates for both paths in Figure 5.

Neutrophils and macrophages generate HOCl in response to foreign insults. The long-standing and widespread rationale for this HOCl production is to serve the singular role of killing invading organisms. Not surprisingly, a substantial amount of time and effort has been dedicated to understanding the killing mechanism therein. Our data provides evidence for an additional role for HOCl by activating the enzyme rhIDO1, a key immune response modulator. Consequently, the results suggest a new signaling pathway in immunomodulation, paving the way for downstream studies to explore the biological implications of this newfound HOCl-IDO1 immunomodulation connection. Just as significant are the mechanistic implications for catalysis. To our knowledge this is the first example of an enzyme that can use HOCl to complete an oxygenation cycle. We hypothesize HOCl is used to generate a Compound I intermediate (both paths) and potentially Compound II (path b), the latter of which has been reportedly observed with Raman spectroscopy and IDO1 prepped in distinct conditions to that reported here (41,42,94). Interestingly, this HOCl-mediated IDO1 catalysis is in contrast to results exploring a potential peroxide shunt using H_2_O_2_, which has led to the conclusion that H_2_O_2_ facilitates the formation of a ferryl species resembling Compound I (50,51) or Compound II (95) yet does not impart IDO1 the ability to transform L-Trp to L-NFK. Furthermore, it is intriguing to speculate that the hypohalite-shunt might not be limited to rhIDO1, with other proximal His- or perhaps proximal Cys-hemoproteins potentially being able to operate in a similar manner (67). Interestingly, LPO in the resting state is already known to convert HOCl to Compound I, in what is essentially the reverse reaction (58). Finally, we are proposing that the ferrous heme is not required, hence exploits the shunt-like pathway in such a manner that O_2_ is not a direct, heme ligand during the catalytic cycle. This last proposal runs counter to the apparently universally accepted paradigm for heme-dependent oxygenases. However, nature has demonstrated multiple other heme-independent strategies to activate O_2_, including oxygenases such as iron-independent flavin-dependent enzymes exemplified by glucose oxidase, ferric-hydroxide-dependent lipoxygenases, and so-called cofactorless oxygenases, among others. It will now be of mechanistic interest to probe the O_2_ activation hypothesis and reconcile this new IDO1 activity with the wealth of data already available in the public domain on this fascinating enzyme.

## METHODS

Materials were from standard sources and used without further purification. The recombinant human IDO1 (rhIDO1) was produced and purified using standard conditions. Routine assays (identified as optimal conditions) were carried out in a 100 mM potassium phosphate buffer (pH = 7.4), 5-10 mM ascorbic acid (AA), 1 mM L-Trp, 0.2-1 mM HOCl, and 0.4 µM rhIDO1. Reactions were performed in an open atmosphere at room temperature. Kinetic data were fitted to the Michaelis-Menten equation using GraphPad Prism 10 to extract parameters. Detailed protein purification protocol, assay conditions including for kinetic analysis, HPLC methods, and production and characterization of *N*-Me-dNFK are provided in the online supporting information.

## ACKNOWLEDGEMENTS

Parts of this work were supported by NIH P20GM130456 COBRE for Translational Chemical Biology.

