## Supporting Information for "The endogenous metabolite hypochlorite activates indoleamine 2,3-dioxygenase-1 for catalysis: Functional and mechanistic implications"

**Table of Contents**

1. **Materials and General Considerations……………………………………………………..S3**
2. **Experimental Methods……………………………………………………………….……S3-S9**
   2.1. Production and purification of rhIDO1………………………………………..…….….......S3
   2.2. Activity assay of rhIDO1 (enzymatic formation of l-NFK and dNFK)…………...……...S5
   2.3. Enzymatic formation of l-NFK and dNFK with hypohalous acids generator system

[LPO + H_2_O_2_ + (pseudo)halides]…………………………………………………………....S6

2.4. Kinetic analysis of rhIDO1 with various activators/ cofactors…………………………...S6
2.5. Total turnover number (TTN) calculation……………………………………………....….S7
2.6. Effect of O_2_ on rhIDO1 reactivity……………………………………………………………S7
2.7. Enzymatic formation of (*E*)-4-[2-(*N*-methylformamido)phenyl]-4-oxo-2-butenoic acid

(*N'*-Me-dNFK)………………………………………………………………………………...S7

2.8. Kinetic analysis of (*E*)-4-(2-(*N*-methylformamido)phenyl)-4-oxo-2-butenoic acid

(*N'*-Me-dNFK) by HPLC……………………………………………………………………..S8

3. **Tables and Figures…………………………………………………………………..…S10-S19**
   3.1. **Table S1.** Activity of rhIDO1 in the presence of variable [H_2_O_2_] and [NaOCl]……….S10
   3.2. **Figure S1:** Effect of pH on rhIDO1 activity in the presence of 200 µM NaOCl in

100 mM potassium phosphate buffers…………………………………………………...S11

3.3. **Figure S2:** Reaction catalyzed by *Canis lupus dingo* indoleamine 2,3-dioxygenase-1 (*cd*IDO1)……………………………………………………………………………………..S12
3.4. **Figure S3:** rhIDO1 catalyzes conversion of 1-Me-l-Trp to *N*-Me-dNFK as a mixture of rotamers……………………………………………………………………………………..S13

3.5. **Figure S4:** Spectroscopic analysis of (*E*)-4-[2-(*N*-methylformamido)phenyl]-4-oxo-2-butenoic acid (*N*-Me-dNFK)…………………………………………………………S14-S18
3.6. **Figure S5:** Effect of O_2_ on rhIDO1 activity……………………………………………….S19
3.7. **Figure S6:** Ozonolysis of l-Trp in aqueous solution……………………………………S19

1. **Materials and General Considerations.**

All chemicals and solvents were purchased from standard commercial sources and used as received without further purification. Ca(OCl)_2_, NaOCl, Br_2_(aq), I_2_, 5-aminolevulinic acid, potassium ferricyanide (III), l-Trp, d-Trp, 1-Me-l-Trp, and LPO from bovine milk (≥200 U/mg) were purchased from Millipore Sigma. Hydrogen peroxide (H_2_O_2_) was purchased as a 30% (w/w) solution from CVS pharmacy (Equate brand) or 3% (w/w) solution with 200 ppm acetanilide stabilizer from Millipore Sigma. The gene encoding human IDO1 was synthesized and cloned in pET22b(+) by GenScript.

HPLC analysis was performed on a Waters
Acquity system equipped with a photodiode array detector and analytical xBridge C18 column (4.6 × 50 mm, 3.5 μm) with a flow rate of 0.4 mL/min. A series of linear gradient was developed from water with 0.1 % formic acid as mobile phase A and acetonitrile with 0.1% formic acid as mobile phase B. The elution program was (beginning time and ending time with linear change to % A) :0 min 98% A; 0-2 min 98% A; 2-20 min 20% A; 20-24 min 20% A; 24-25 min 98% A; 25-35 min 98% A.  High-resolution mass spectrometry (HR-ESI-MS) data were acquired with an Agilent 6230 TOF LC/MS System (Agilent Technologies) equipped with a Gemini 11CA C18 column (4.6 × 100 mm, 3 μm) with a flow rate of 0.4 mL/min. A series of linear gradients were developed from water with 0.1% formic acid as mobile phase A and acetonitrile with 0.1% formic acid as mobile phase B. The elution program is identical to the above.  The UV/VIS data was collected using a Shimadzu TCC-240A or a Thermo Scientific NanoDrop™ ND-1000 spectrometer.

2. **Experimental Methods.**
   2.1. **Production and purification of rhIDO1.**

Plasmids pET22b(+)-*ido1* was introduced into *E. Coli* BL21(DE3) cells, which were then plated on LB (Luria-Bertani) agar supplemented with 50 μg/mL of  ampicillin and incubated overnight at 37 °C. A single colony was used to inoculate 5 mL of TB (Terrific Broth) or LB (Luria-Bertani) medium with 50 μg/mL of ampicillin, and the culture was shaken at 37 °C for 8 hours. Subsequently, 500 µL of this culture was used to inoculate 50 mL of TB or LB medium supplemented with 50 μg/mL of ampicillin and the flask was incubated on a shaker overnight at 37 °C. Next, 10 mL of the 50 mL culture was transferred to 500 mL of TB or LB medium supplemented with 50 μg/mL of ampicillin and incubated at 18 °C with shaking until an OD of ~0.5 was reached. Approximately one hour before induction, 500 μL of 50 mg/ mL 5-aminolevulinic acid solution was added to each flask. Protein expression was induced by the addition of isopropyl *b*-D-1-thiogalactopyranoside (IPTG) to a final concentration of 100 μM, and cells were shaken overnight at 18 °C before being harvested by centrifugation.

The cell paste was resuspended in the lysis buffer containing 100 mM potassium phosphate buffer (pH = 7.4), 200 mM KCl, and 5 mM imidazole. Cell lysis was performed using Qsonica sonicator (Qsonica LLC, Newtown, CT) at 65-69% amplitude for 5 cycles. Each cycle is separated by 30 seconds of rest time and consists of 1-second pulse separated by 1-second rest periods for the total time of 30 seconds. Cell debris was then removed by centrifugation, and the supernatant was loaded onto a Ni-NTA agarose column (Thermo Scientific, Rockford, IL) to purify the recombinant proteins using 6XHis-tag at the C-terminus. The column was washed three times with 10 mL of wash buffer containing 100 mM potassium phosphate buffer (pH = 7.4), 200 mM KCl, and 25 mM imidazole. The target proteins were eluted with 100 mM potassium phosphate buffer (pH = 7.4) and 250 mM imidazole. Purified proteins were concentrated and desalted using 100 mM potassium phosphate buffer (pH = 7.4) and a PD-10 column, then flash frozen and stored at –80 °C. Additionally, a portion of the concentrated enzyme was incubated on ice for 20 min after the addition of 1 mM potassium ferricyanide (III) solution, prior to desalting with a PD-10 column, to ensure the formation of ferric (Fe^+3^)-rhIDO1. (Note: Ferricyanide-treated rhIDO1 (FC-rhIDO1) was only tested with Ca(OCl)_2_, NaOCl, and I_2_/NaOH activators). The enzyme purity was assessed by SDS-PAGE followed by Coomassie blue staining and appeared as a dominant band around 46 kDa. Protein concentration was determined by UV/VIS spectroscopy (ε_404_ = 172 mM^-1^cm^-1^ for rhIDO1) using NanoDrop™ ND-1000 spectrometer.

rhIDO1 sequence:

MAHAMENSWTISKEYHIDEEVGFALPNPQENLPDFYNDWMFIAKHLPDLIESGQLRERVEKLNMLSIDHLTDHKSQRLARLVLGCITMAYVWGKGHGDVRKVLPRNIAVPYCQLSKKLELPPILVYADCVLANWKKKDPNKPLTYENMDVLFSFRDGDCSKGFFLVSLLVEIAAASAIKVIPTVFKAMQMQERDTLLKALLEIASCLEKALQVFHQIHDHVNPKAFFSVLRIYLSGWKGNPQLSDGLVYEGFWEDPKEFAGGSAGQSSVFQCFDVLLGIQQTAGGGHAAQFLQDMRRYMPPAHRNFLCSLESNPSVREFVLSKGDAGLREAYDACVKALVSLRSYHLQIVTKYILIPASQQPKENKTSEDPSKLEAKGTGGTDLMNFLKTVRSTTEKSLLKEGLEHHHHHH

2.2. **Activity assay of rhIDO1 (enzymatic formation of l-NFK and dNFK).**

Reactions were carried out in a 100 mM potassium phosphate buffer (pH = 7.4), containing 5-10 mM ascorbic acid (AA), 1 mM l-Trp, various concentration of Activators/ cofactors [Ca(OCl)_2_, NaOCl, I_2_/ NaOH, Br_2_/ H_2_O, NaOCl + NaI_2_, NaOCl + NaBr, and methylene blue], and 0.4 µM rhIDO1. The mixtures were incubated at RT for 2.5 h and 18 h. After incubation, the mixtures were boiled for 2 minutes, cooled down on ice, and centrifuged to remove the protein at 14000 rpm for 10 minutes. The supernatant was analyzed by HPLC. The products concentrations were determined by integrating peak areas (A) and applying Beer-Lambert’s law, using the absorption coefficients ε_280_ = 3075 M^-1^cm^-1^ for dNFK and ε_321_ = 3750 M^-1^cm^-1^ for l-NFK. The remaining substrate concentration was calculated using an absorption coefficient of ε_280_ = 5690 M^-1^cm^-1^. The relative percent conversion was calculated and plotted for each cofactor/ activator.

| **Activator/Cofactor** | **[Activator/ cofactor]** | **[l-Trp]** | **[AA]** | **Time** |
| --- | --- | --- | --- | --- |
| NaOCl | 0 µM – 1 mM | 1 mM | 0 mM – 10 mM | 0 – 18 h |
| Ca(OCl)_2_ | 0 µM – 1 mM | 1 mM | 0 mM – 10 mM | 0 – 18 h |
| I_2_/ NaOH | 0 µM – 5 mM | 1 mM | 0 mM – 10 mM | 0 – 18 h |
| Br_2_/H_2_O | 0 µM – 5 mM | 1 mM | 0 mM – 10 mM | 0 – 18 h |
| NaOCl +NaI | NaOCl (0 µM – 1 mM)  NaI (0 µM – 10 mM) | 1 mM | 10 mM | 0 – 18 h |
| NaOCl +NaBr | NaOCl (0 µM – 1 mM)  NaBr (0 µM – 10 mM) | 1 mM | 10 mM | 0 – 18 h |
| Methylene blue | 0 µM – 10 µM | 1 mM | 0 mM – 10 mM | 0 – 18 h |

2.3. **Enzymatic formation of l-NFK and dNFK with hypohalous acids generator system [LPO + H_2_O_2_ + (pseudo)halides]**

Reactions were prepared in 100 mM potassium phosphate buffer (pH = 7.4), containing 1 mM l-Trp, 50 mM (pseudo)halide salts (KF, KCl, KBr, KI, NaBr, NaI, and KSCN), 20 µg/mL of lactoperoxidase, and 0.5 µM rhIDO1. Reactions were initiated by the addition of 2 mM H_2_O_2_ and carried out in the absence or presence of 10 mM AA for 3 h at RT. After incubation, the reactions were worked up by heat for 2 minutes, cooled down on ice, and centrifuged to remove the protein at 14000 rpm for 10 minutes. The supernatant was analyzed by HPLC. The products concentrations were determined by integrating peak areas (A) and applying Beer-Lambert’s law, using the absorption coefficients ε_280_ = 3075 M^-1^cm^-1^ and ε_321_ = 3750 M^-1^cm^-1^ for dNFK and l-NFK, respectively. The remaining substrate concentration was calculated using an absorption coefficient of ε_280_ = 5690 M^-1^cm^-1^. The percent conversions were then calculated and plotted versus various concentrations of each salt.

2.4. **Kinetic analysis of rhIDO1 with various activators/ cofactors.**

The kinetic parameters for rhIDO1 in the presence of each cofactor/ activator were determined at RT in a quartz cuvette using UV/VIS spectrometer. The reaction mixtures (100 µL) contained 100 mM potassium phosphate (pH = 7.4), 10 mM ascorbic acid (AA), varying concentrations of l-Trp (50 µM – 3.2 mM), and 0.2 µM rhIDO1. The reactions were initiated by adding each cofactor/ activator, and the increase in product (l-NFK) absorbance was monitored over time at λ_max_ = 321 nm. Initial velocities were calculated under the steady-state condition considering the linear region of the curve. Each data point represents at least two replicates, and Michaelis–Menton kinetic constants were determined by nonlinear regression analysis using GraphPad Prism 10 (GraphPad Software, La Jolla, CA, USA). The kinetic parameters are summarized in Figure 4 and Table 2.

2.5. **Total turnover number (TTN) calculation.**

Total turnover numbers (TTNs) were determined at RT with a quartz cuvette using UV/VIS spectrometer. In a cuvette, the reaction mixture contained 100 mM potassium phosphate (pH = 7.4), 10 mM ascorbic acid, 1 mM l-Trp, and ~0.05 µM of rhIDO1. Reactions were initiated by addition of 100 µM or 500 µM Ca(OCl)_2_. The formation of l-NFK was monitored in real time by measuring absorbance at λ_max_ = 321 nm. The Maximum absorbance was recorded when rhIDO1 activity ceased, and the product signal reached a plateau. l-NFK concentrations were calculated from the maximum absorbance values using the Beer-Lambert’s law, and TTNs were determined using the following formula:

$$TTN= \frac{Moles of the product}{Moles of enzyme}$$

2.6. **Effect of O_2_ on rhIDO1 reactivity.**

In a two-neck pear-shaped flask, 500 µL of 100 mM potassium phosphate buffer (pH = 7.4) and 1 mM l-Trp were prepared. In a separate flask, 500 µL of the same buffer and 5 µM of IDO1 were combined. Both flasks were placed under an N_2_ flow using a Schlenk line, and the mixtures inside were directly purged with the gentle flow of N_2_ gas via needles for 1.5 h to remove oxygen. Then, iodine (0.5 mg, 2 µmol) was added to the first flask solution, sealed, and gently sonicated until dissolved. The enzyme solution was then transferred to the first flask using a cannula, and the reaction mixture was kept under the N_2_ flow for about 4 h. For the control reaction, the mixture was exposed to air by removing the cap and discontinuing the N_2_ flow after addition of the enzyme solution. Reaction progress for both the control and experimental condition, repeated in duplicate, was monitored using HPLC.

2.7. **Enzymatic formation of (*E*)-4-[2-(*N*-methylformamido)phenyl]-4-oxo-2-butenoic acid** **(*N*-Me-dNFK)**

Reaction consisted of 100 mM potassium phosphate buffer (pH = 7.4), 1 mM 1-Me-l-Trp, 10 mM AA, and 0.4 µM rhIDO1 was initiated by addition of 200 µM NaOCl. The reaction was performed at RT and subjected to heat work-up for 2 minutes after 18 h. The enzyme was removed by centrifugation at 14000 rpm for 10 minutes and the product formation was analyzed by HPLC and purified as a mixture of rotamers (2:1 ratio) by preparative HPLC using SunFire Prep C18 column (5 µm, 10 × 250 mm) at flow rate of 3.5 mL/min. A series of linear gradients was developed from water with 0.1% formic acid as mobile phase A and acetonitrile with 0.1% formic acid as mobile phase B. The elution program was: 0 min, 2% B; 0-2 min 2% B; 2-14 min, 80% B; 14-16 min, 80% B; 16-17 min, 2% B; 17-19 min, 2% B. The elution was monitored at 280 nm. ^1^H NMR (600MHZ, CD_3_OD) 8.14 (s, 0.5H, minor), 8.07 (s, 1H, major), 7.77 (d, *J* = 7.8 Hz, 1H, major), 7.71 (t, *J* = 7.8 Hz, 1H, major), 7.69 (d, *J* = 7.8 Hz, 0.5H, minor), 7.68 (t, *J* = 7.8 Hz, 0.5H, minor), 7.56 (t, *J* = 7.8 Hz, 1H, major), 7.51 (t, *J* = 7.8 Hz, 0.5H), 7.48 (d, *J* = 7.8 Hz, 1H, major), 7.41 (d, *J* = 7.8 Hz, 0.5H, minor), ,7.38 (d, *J* = 16.0 Hz, 1H, major), 7.36 (d, *J* = 15.8 Hz, 0.5H, minor), 6.62 (d, *J* = 16.0 Hz, 1H, major), 6.61 (d, *J* = 15.8 Hz, 0.5H, minor), , 3.43 (s, 1.5H, minor), 3.23 (s, 3H, major). ^13^C NMR (150 MHz, CD_3_OD) 193.7, 193.4, 168.1, 165.2, 164.9, 142.0, 140.0, 139.5, 137.2, 136.5, 135.0, 134.7, 134.2, 131.3, 130.3, 129.4, 129.2, 129.0, 128.3, 38.2, 34.3. HRMS (ESI) Calcd. for C_12_H_12_NO_4_ [M+H]^+^: 234.0761; Found: 234.0755.

2.8. **Kinetic analysis of (*E*)-4-(2-(*N*-methylformamido)phenyl)-4-oxo-2-butenoic acid (*N*-Me-dNFK) by HPLC**

Enzymatic reactions were performed at varying concentration of 1-Me-l-Trp (50 µM – 2.4 mM) in 100 mM potassium phosphate (pH = 7.4), and 10 mM ascorbic acid (AA). Reactions were initiated by addition of 0.2 µM of rhIDO1 and 200 µM NaOCl, respectively and allowed to proceed under initial rate conditions. At specific time intervals (2, 4, 6, 8,10, and 15 min), reactions were quenched by boiling for 2 minutes, then cooled down on ice, and centrifuged to remove the protein at 14000 rpm for 10 minutes. Samples were analyzed by HPLC, and product formation was quantified based on calibration curves. The absorption coefficients were determined to be ε_222_ = 34430 M^-1^cm^-1^ and ε_227_ = 14810 M^-1^cm^-1^ for 1-Me-l-Trp and the product, respectively. Initial velocities were calculated under the steady-state condition from the linear range of product formation versus time plots. Each data point represents at least two replicates, and Michaelis–Menton kinetic constants were determined by nonlinear regression analysis using GraphPad Prism 10 (GraphPad Software, La Jolla, CA, USA). The kinetic parameters are summarized in Figure 4 and Table 2.

**Table S1. Activity of rhIDO1 in the presence of variable [H_2_O_2_] and [NaOCl]^a^**

| Activator/Cofactor | 0 mM H_2_O_2_ | 1 mM H_2_O_2_*^b^* | 2 mM H_2_O_2_*^b^* | 10 mM H_2_O_2_ |
| --- | --- | --- | --- | --- |
| 0 µM NaOCl*^c^* | – | NR*^e^* | NR | NR |
| 0 µM NaOCl*^d^* | – | NR | NR | NR |
| 200 µM NaOCl | 29.7*^f^* | 32.2 | 26.9 | 6.5 |
| 1 mM NaOCl | 48.6 | 63.4 | 54.6 | 21.4 |

*^a^*Reaction condition: 100 mM potassium phosphate (pH = 7.4), 0-10 mM AA, 1 mM l-Trp, 0-10 mM H_2_O_2_, and 0.4 µM rhIDO1; *^b^*The slight increase in the %conversion at 1-2 mM H_2_O_2_ is likely due to the in situ generation of O_2_(g) from the reaction between H_2_O_2_ and NaOCl (NaOCl + H_2_O_2_ ⇋ NaCl +H_2_O + O_2_); *^c^*with 10 mM AA; *^d^*without AA; *^e^*No Reaction; *^f^*each value represents the calculated % conversion.

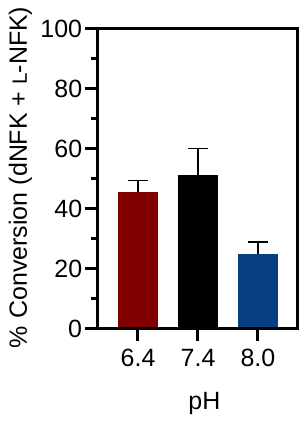

**Figure S1: Effect of pH on rhIDO1 activity in the presence of 200 µM NaOCl in 100 mM potassium phosphate buffers.** The optimal pH was determined to be 7.4, consistent with physiological conditions. Products formation (l-NFK + dNFK) at pH 6.4 was comparable to that observed at pH = 7.4, with the higher amount of l-NFK formed. Increasing the pH to 8.0 resulted in an approximately 50% decrease in percent conversion, with only trace amount of l-NFK formed.

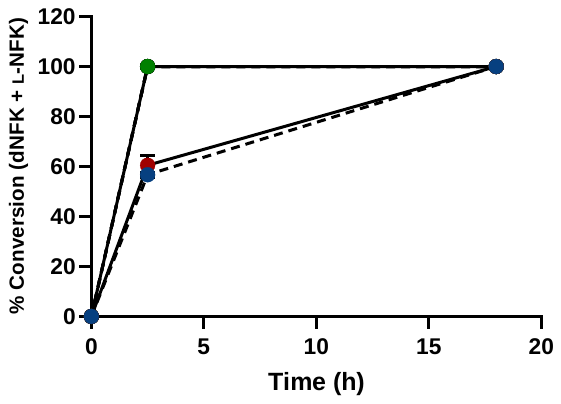

**Figure S2: Reaction catalyzed by Canis lupus dingo indoleamine 2,3-dioxygenase-1 (cdIDO1).** Reactions were monitored for 2.5 h and 18 h with 10 mM ascorbic acid and 100 µM Ca(OCl)_2_ ( ● ); 200 µM NaOCl ( ● ); and 500 µM Ca(OCl)_2_ or 1mM NaOCl or 1mM I_2_/ NaOH or 1 mM Br_2_/ H_2_O ( ● ).

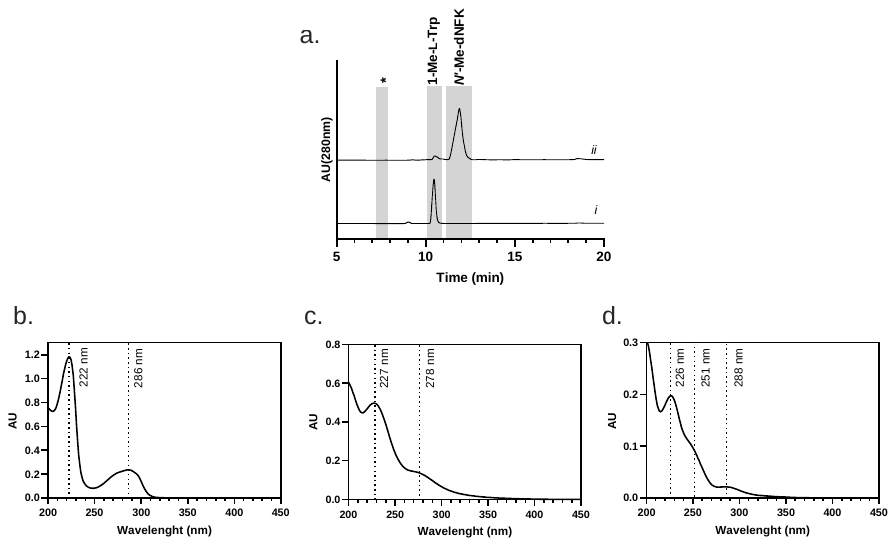

**Figure S3: rhIDO1 catalyzes conversion of 1-Me-l-Trp to N-Me-dNFK as a mixture of rotamers.** (a) HPLC traces of the rhIDO1-catalyzed reaction including the control (no enzyme, trace i); rhIDO1 reaction using AA and NaOCl followed by heat work up (trace ii); *, retention time for N'-Me-l-NFK which is not observed at 280 nm, as it is completely converted to N-Me-dNFK following the heat work up. The retention time for N'-Me-l-NFK was determined after processing the sample obtained following the work up by ultrafiltration; (b) UV/VIS spectrum of 1-Me-l-Trp; (c) UV/VIS spectrum of N-Me-dNFK; (d) UV/VIS spectrum of N'-Me-l-NFK.

a.

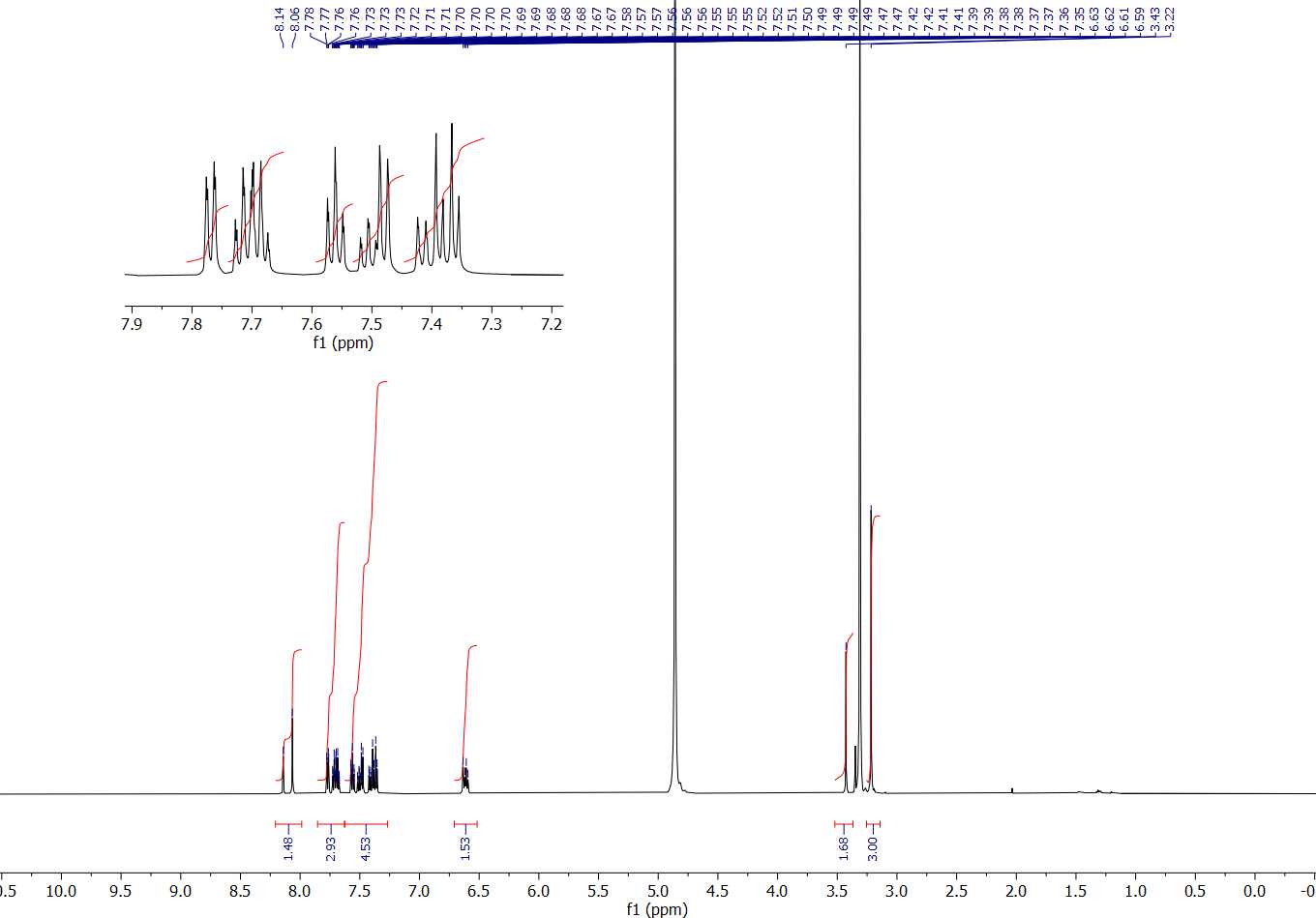

b.

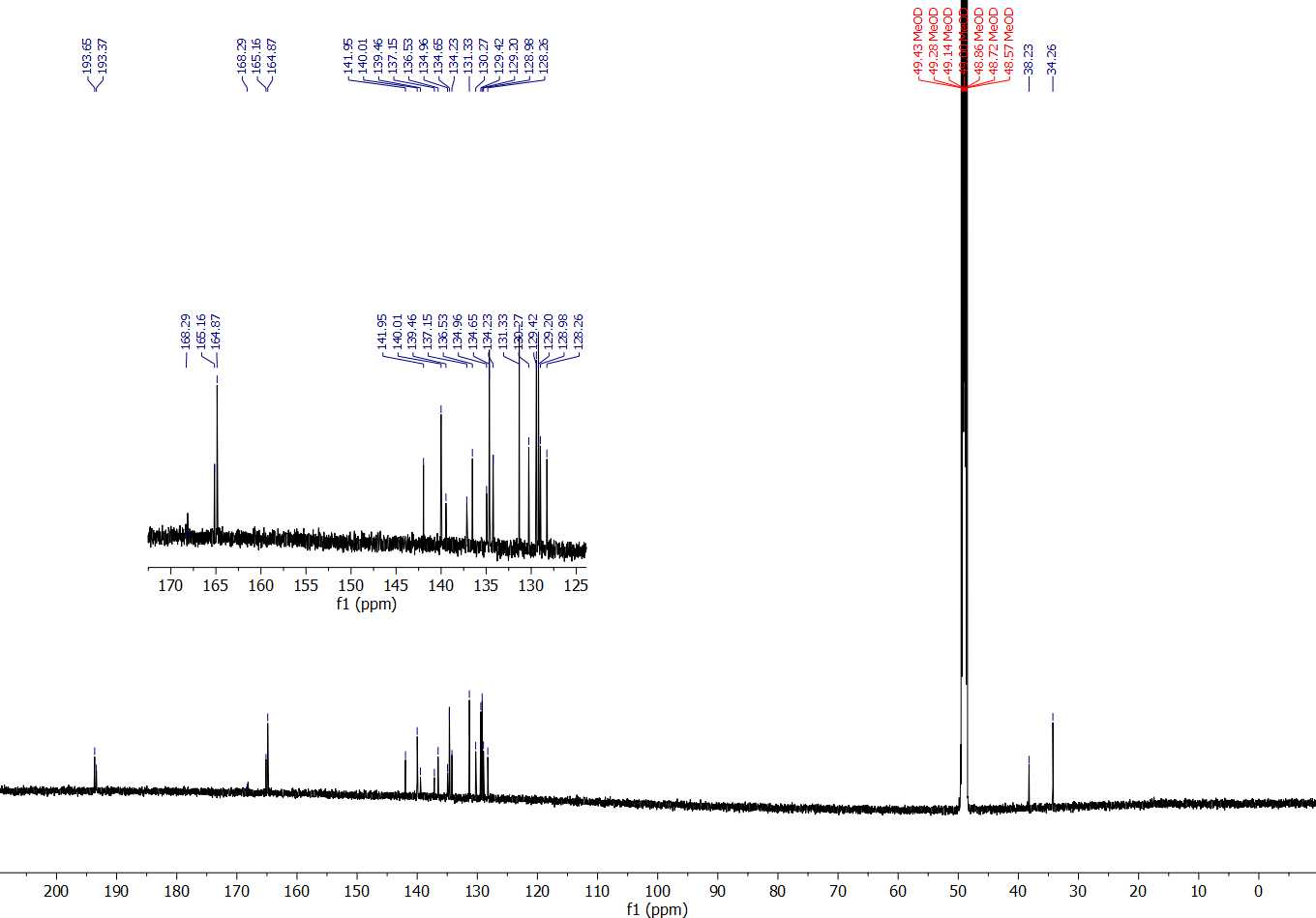

c.

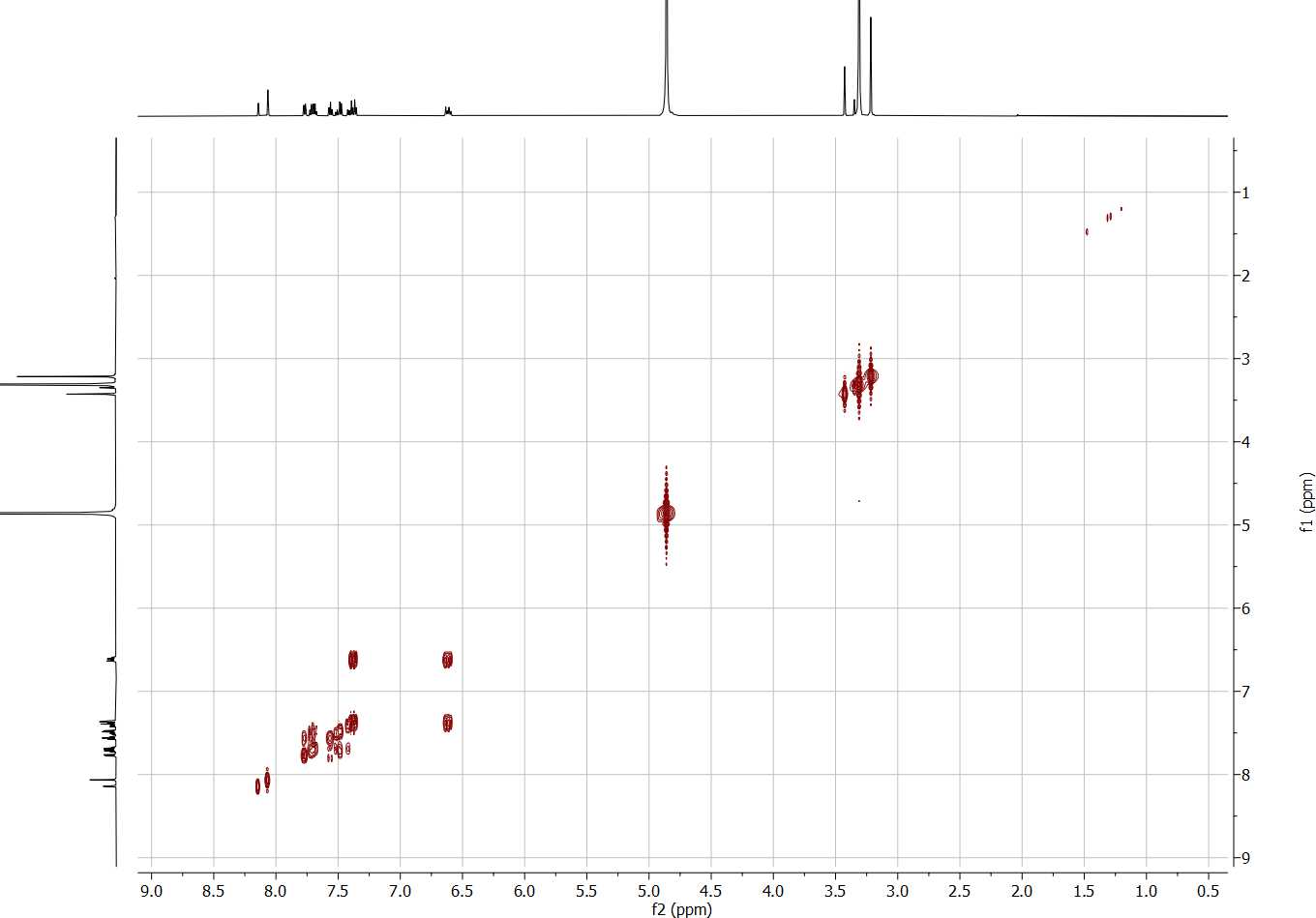

d.

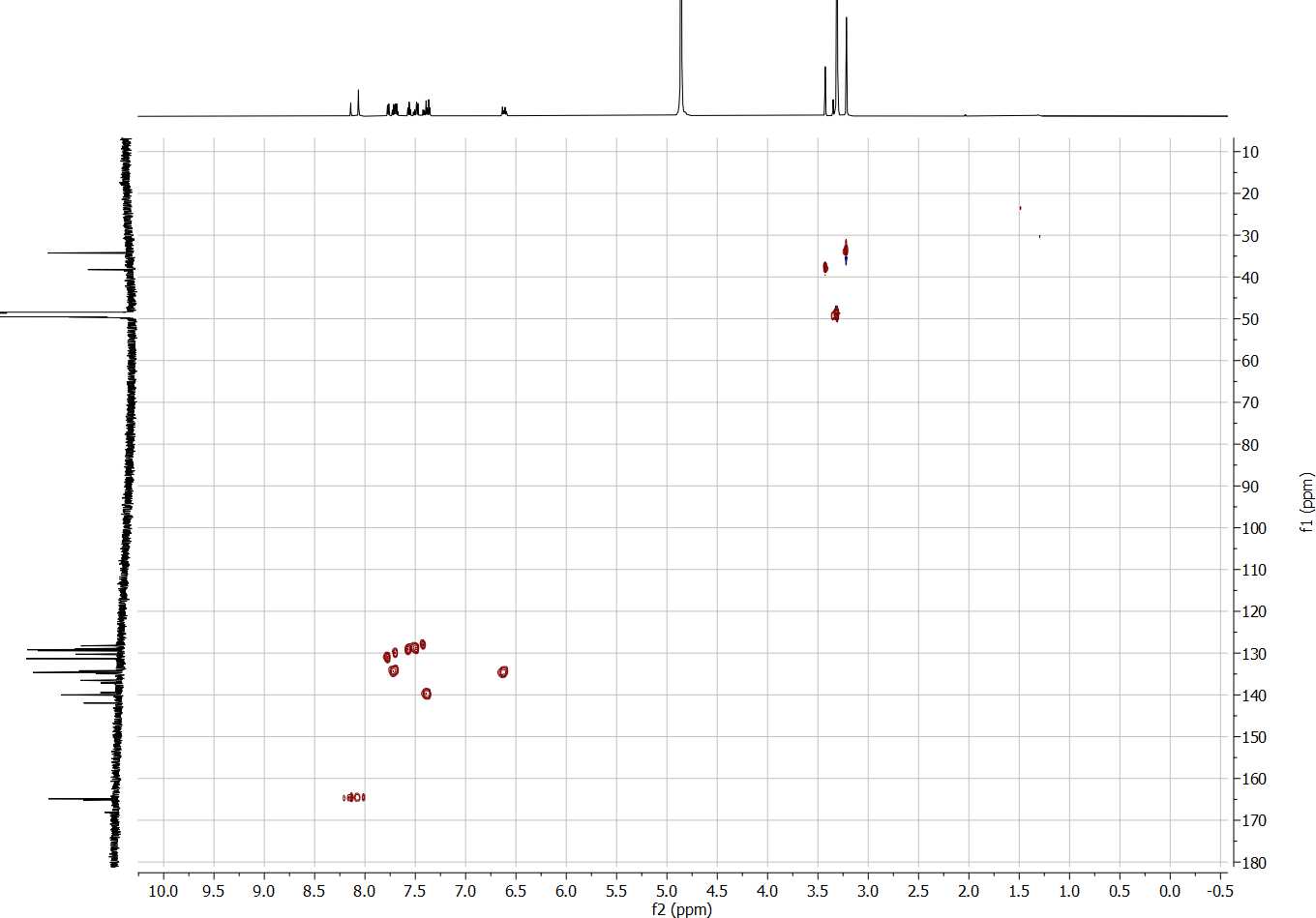

e.

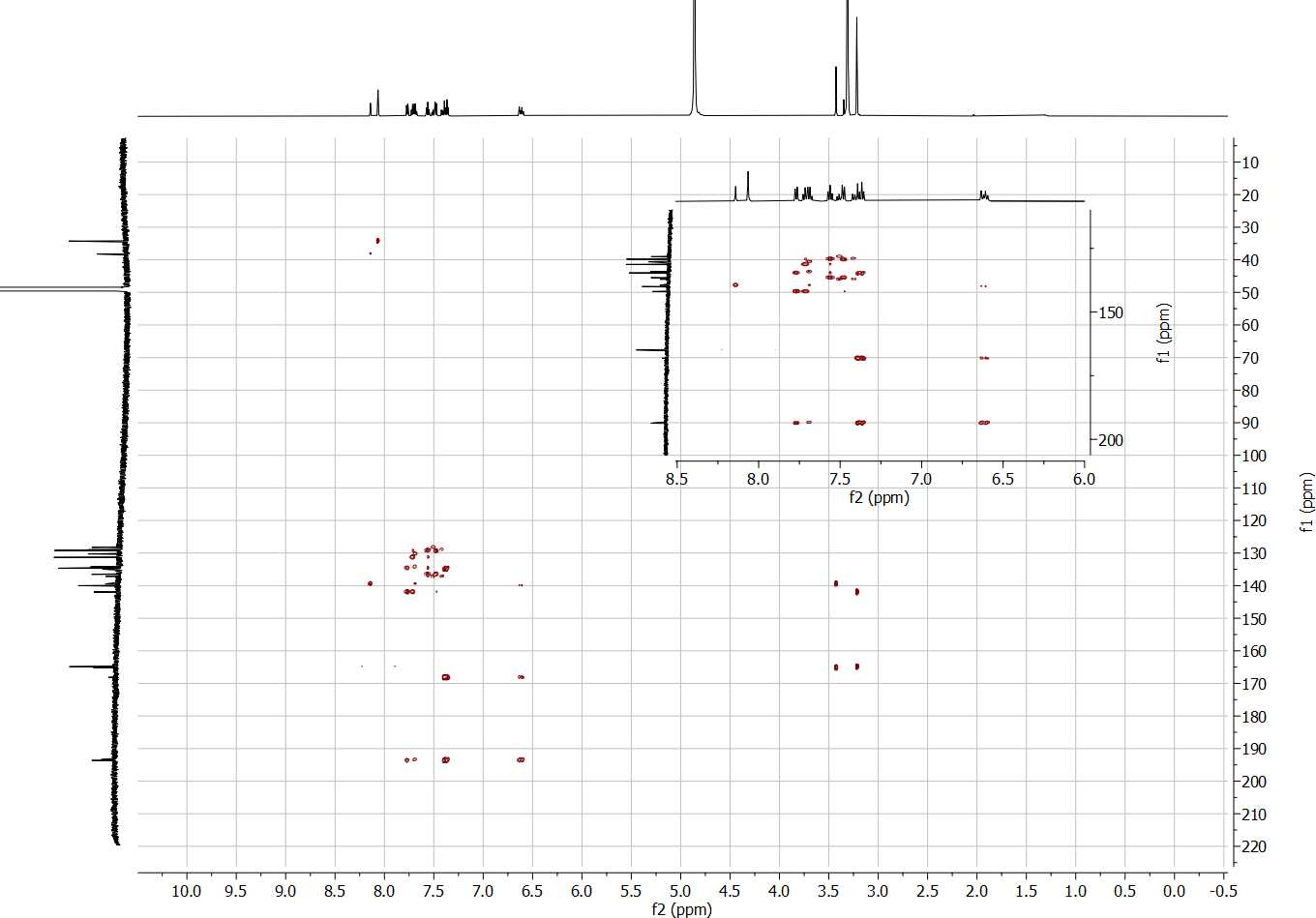

f.

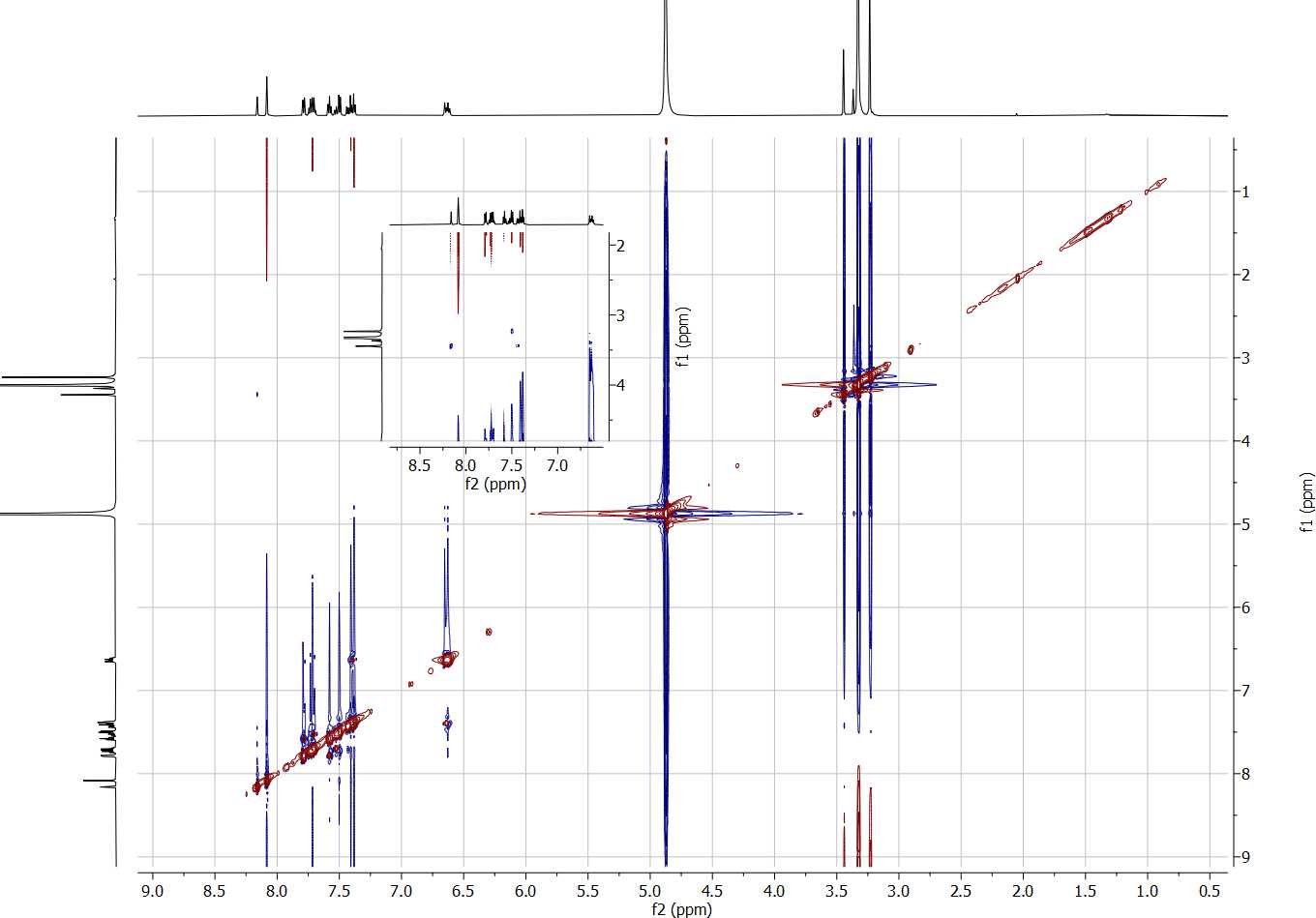

g.

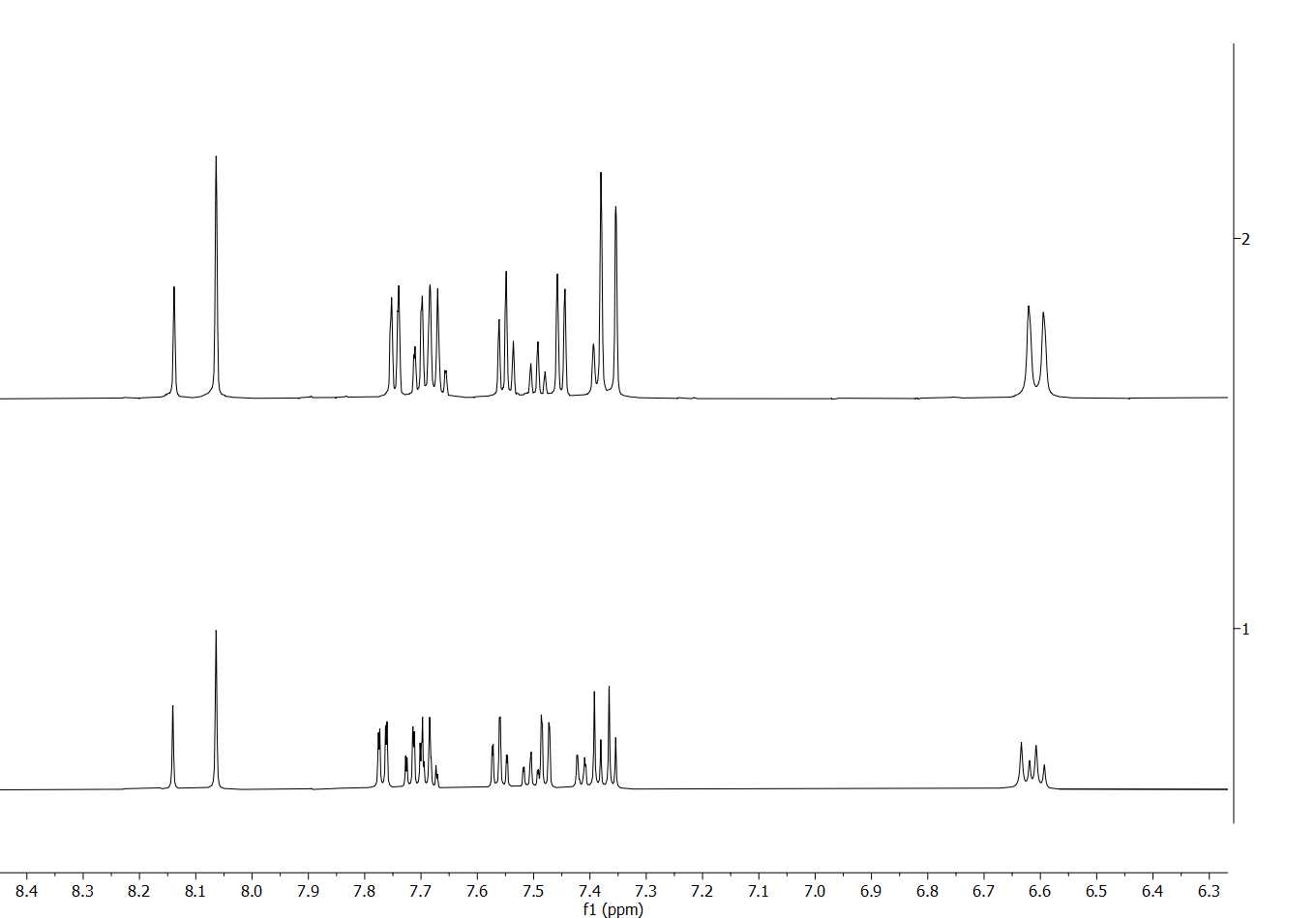

**25 °C**

**55 °C**

h.

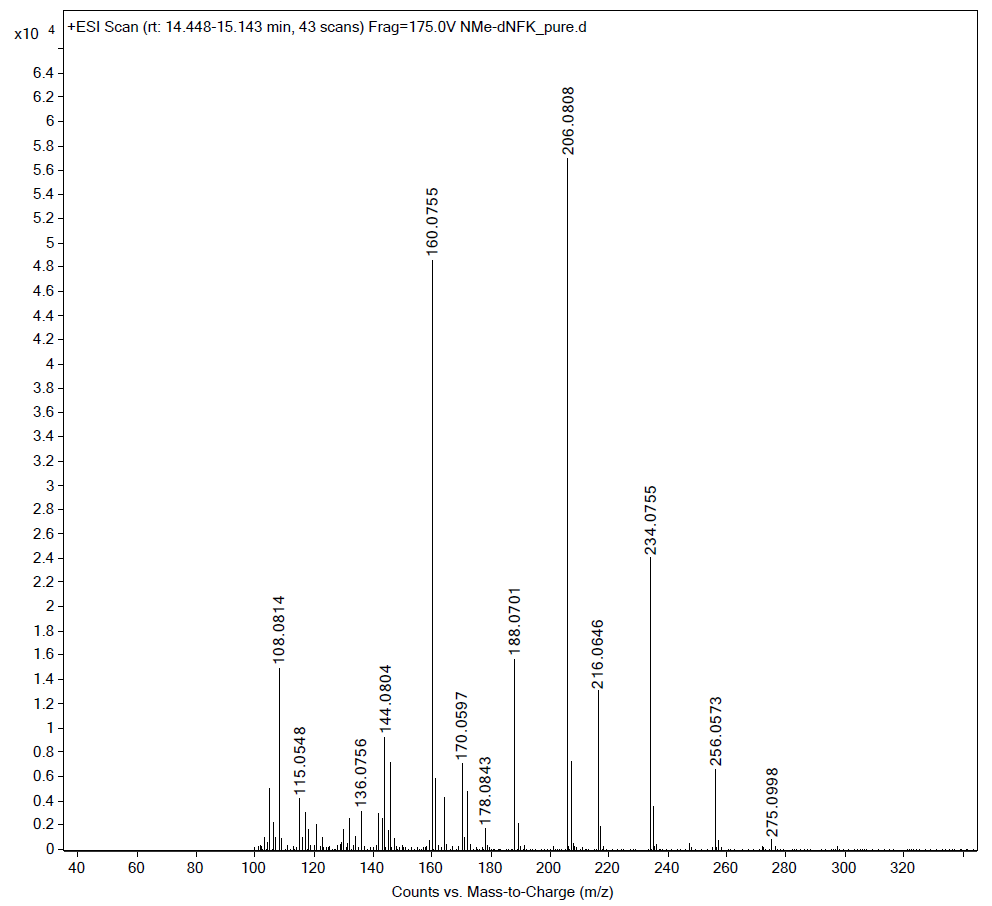

**[M+H]^+^**

**[M+Na]^+^**

**Figure S4:** **Spectroscopic analysis of (*E*)-4-[2-(*N*-methylformamido)phenyl]-4-oxo-2-butenoic acid (*****N*-Me-dNFK).** a)^1^H NMR spectrum (CD_3_OD, 600 MHz); b)^13^C NMR spectrum (CD_3_OD, 150 MHz); c) ^1^H-^1^H COSY (CD_3_OD, 600 MHz); d) ^1^H-^13^C HSQC (CD_3_OD, 600 MHz); e) ^1^H-^13^C HMBC (CD_3_OD, 600 MHz); f) ^1^H-^1^H NOESY (CD_3_OD, 600 MHz); g) Stacked ^1^H NMR spectra at 55 °C (top) and RT (bottom). Faster rotation at higher temperature was achieved, leading to coalescence of the rotamers, which is observable in the alkene protons at ~6.60 and ~7.35 ppm. h) (+)-HR-ESI-MS (expected [M+H]^+^ at *m/z* = 234.0761, found 234.0755)

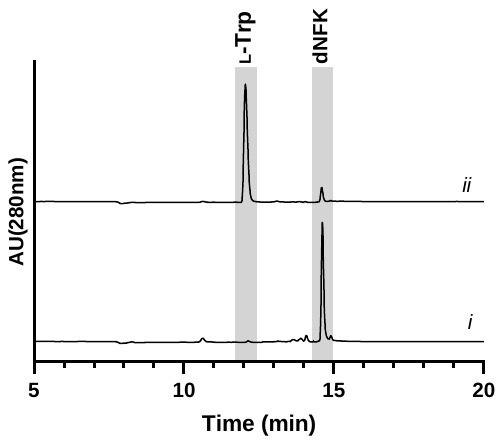

**Figure S5: Effect of** **O_2_ on rhIDO1 activity**. rhIDO1 catalyzes complete conversion of l-Trp to dNFK in the presence of AA and I_2_/NaOH under an O_2_ atmosphere (trace i), whereas it produces ~7% (by HPLC) of dNFK under an anaerobic condition (trace ii). HPLC analysis was performed at 280 nm following the heat work-up.

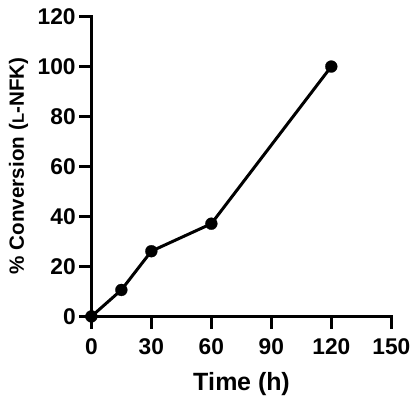

**Figure S6:** **Ozonolysis of l-Trp in aqueous solution.** Ozone generator converts 1 mM l-Trp in 100 mM potassium phosphate buffer (pH = 7.4) with 10 mM AA to l-NFK.
